# Leveraging a large harmonized epigenomic data collection for metadata prediction to validate and augment over 350,000 public epigenomic datasets

**DOI:** 10.1101/2025.09.04.670545

**Authors:** Joanny Raby, Gabriella Frosi, Frédérique White, Jonathan Laperle, Pierre-Étienne Jacques

## Abstract

Epigenomic data found in public databases often suffer from issues of non-standardization and incompleteness in their associated metadata. There are currently no automated approaches to validate or correct such missing or inaccurate metadata information. To tackle this challenge, we harnessed the extensive harmonized data and metadata provided by the EpiATLAS project of the International Human Epigenome Consortium (IHEC) to train EpiClass, a suite of machine learning classifiers that can predict key metadata (∼98% accuracy), including experimental assay, donor sex, biospecimen source and sample cancer status. The development of these classifiers enabled the identification of a few mislabeled and low-quality datasets in the EpiATLAS project, while also completing with high-confidence most of the missing metadata. These classifiers were also validated on thousands of ENCODE datasets absent from the initial training, then applied to assess more than 350,000 human ChIP-seq and RNA-seq datasets from public repositories. Overall, this effort not only validated the accuracy of the vast majority of assays reported by the original authors, but also unveiled ∼500 datasets with discrepancies, in particular through data swap during data entry. More importantly, EpiClass also supplied high-confidence predictions for over 320,000 metadata attributes of the biological sample such as the sex, cancer status and biomaterial type, which had been originally omitted in the majority of cases. Our work introduces the first systematic approach for metadata correction and augmentation, enhancing the quality and reliability of publicly available epigenomic data.

## Introduction

The advancement of high-throughput sequencing technologies has driven an exponential increase in publicly available epigenomic data. These data are housed in repositories like the Gene Expression Omnibus (GEO) and the Sequence Read Archive^1,2^, offering unprecedented opportunities to understand gene regulation, cellular identity, and disease mechanisms. Integrating and analyzing this wealth of information, encompassing diverse epigenomic assays such as Chromatin Immuno-Precipitation followed by sequencing (ChIP-seq) for protein localization, transcriptomics (RNA-seq) for gene expression, open chromatin sequencing (e.g., DNase-seq, ATAC-seq) and DNA methylation profiling (e.g., Whole-Genome Bisulfite Sequencing (WGBS)), holds immense promise for accelerating biological discovery and enabling predictive medicine. This potential is highlighted by the use of machine learning approaches to predict some epigenomic assays from others^3–5^, and the growing field of multimodal data analysis for applications such as patient survival and therapy response prediction^6^. Overall, GEO developers report that more than 50,000 publications cite data they host to support independent studies or as the basis of analytical hypotheses and tools^1,7^.

However, harnessing the full potential of these public resources is significantly impeded by pervasive issues with their associated metadata limiting their reusability. Widespread challenges in data reporting and curation practices create critical obstacles for the secondary utilization of these valuable datasets, hindering the application of advanced data science and artificial intelligence approaches^8^. Crucial experimental details, including assay types, biological sources such as cell and tissue type (hereafter termed biospecimen source), donor characteristics, and protocols, are frequently documented inconsistently, incompletely, or within unstructured free-text descriptions^9–11^. This common lack of standardized, machine-readable metadata severely hinders data discovery, large-scale integration, comparative analyses, and the development of robust computational models, ultimately limiting the reliability and scientific value derived from public data repositories

Several computational strategies have been developed to mitigate these metadata challenges. One major line of work focuses on processing the available textual descriptions using techniques ranging from text mining and rule-based systems to machine learning approaches (including large language models) for automatically extracting structured attributes or improving data organization^12–15^. While powerful, the success of these methods is inherently limited by the presence, completeness, and clarity of the original text annotations. A complementary approach that is becoming more relevant involves predicting biological sample characteristics or experimental variables directly from the molecular data itself using machine learning. This has shown particular promise for predicting attributes like donor sex, age, or tissue/cancer type with high accuracy from RNA-seq gene expression profiles or other epigenomic signals like DNA methylation^16–20^. Machine learning has also enabled inference of cellular composition from bulk tissue data^21–23^. These methods collectively demonstrate the potential to infer metadata directly from the data, overcoming gaps in textual descriptions. Other initiatives such as ChIP-Atlas^24^, Cistrome Data Browser^25^, and Next Generation Sequencing Quality Control Generator (NGS-QC)^26^ also reprocessed the raw data in addition to standardizing the metadata from GEO. However, these existing approaches are typically designed for specific assay types or prediction tasks and often rely on predefined genomic features such as gene expression profiles or known regulatory regions. A general approach capable of both validating existing metadata and predicting a broad spectrum of key attributes across diverse epigenomic data types and using whole-genome signal directly without prior feature selection has remained a critical need.

To address this gap, we developed EpiClass, a suite of machine learning classifiers trained on the largest, most diverse and highest-quality collection of harmonized epigenomic datasets, provided by the EpiATLAS project of the International Human Epigenome Consortium (IHEC)^27^. EpiClass is based on the hypothesis that intrinsic signal patterns within different epigenomic assays can robustly predict not only the assay type, but also key characteristics of the biological sample such as biospecimen source, biomaterial type, donor sex, life stage, and sample cancer status. Our models indeed achieved high accuracy across multiple metadata categories and identified mislabeled and low-quality datasets within EpiATLAS itself, in addition to assigning confident labels to missing characteristics. We further validated the models on external ENCODE datasets, motivating their application to over 350,000 public human ChIP-seq and RNA-seq datasets. This large-scale analysis confirmed the majority of existing annotations, while uncovering numerous discrepancies and filling critical gaps with high-confidence predictions such as hundreds of thousands of donor sex and sample cancer status. This work establishes a proof-of-concept providing the first systematic, data-driven method to validate and enrich public epigenomic resources directly from their epigenomic signal profiles, significantly enhancing their quality, reliability, and utility.

## Results

### EpiClass accurately predicts EpiATLAS assay and biospecimen source metadata

The harmonized EpiATLAS data and metadata used to develop EpiClass comprise 7,464 datasets (independent experiments) from 2,216 epigenomes (biological samples) generated by nine international consortia such as ENCODE, Blueprint and CEEHRC. Each epigenome included data from up to nine different assays, which we define as ‘core assays’ for this study: six ChIP-seq histone modifications sharing a single control Input file, RNA-seq and WGBS (Fig. 1a, Supplementary Table 1). The total training set of EpiClass included 20,922 signal files, comprising multiple normalization outputs per ChIP-seq dataset (raw, fold change and p-value track types) and strand-specific files for RNA-seq and WGBS assays. We included multiple track types and stranded tracks to increase classifier robustness to external datasets and expand the training set.

**Figure 1.**
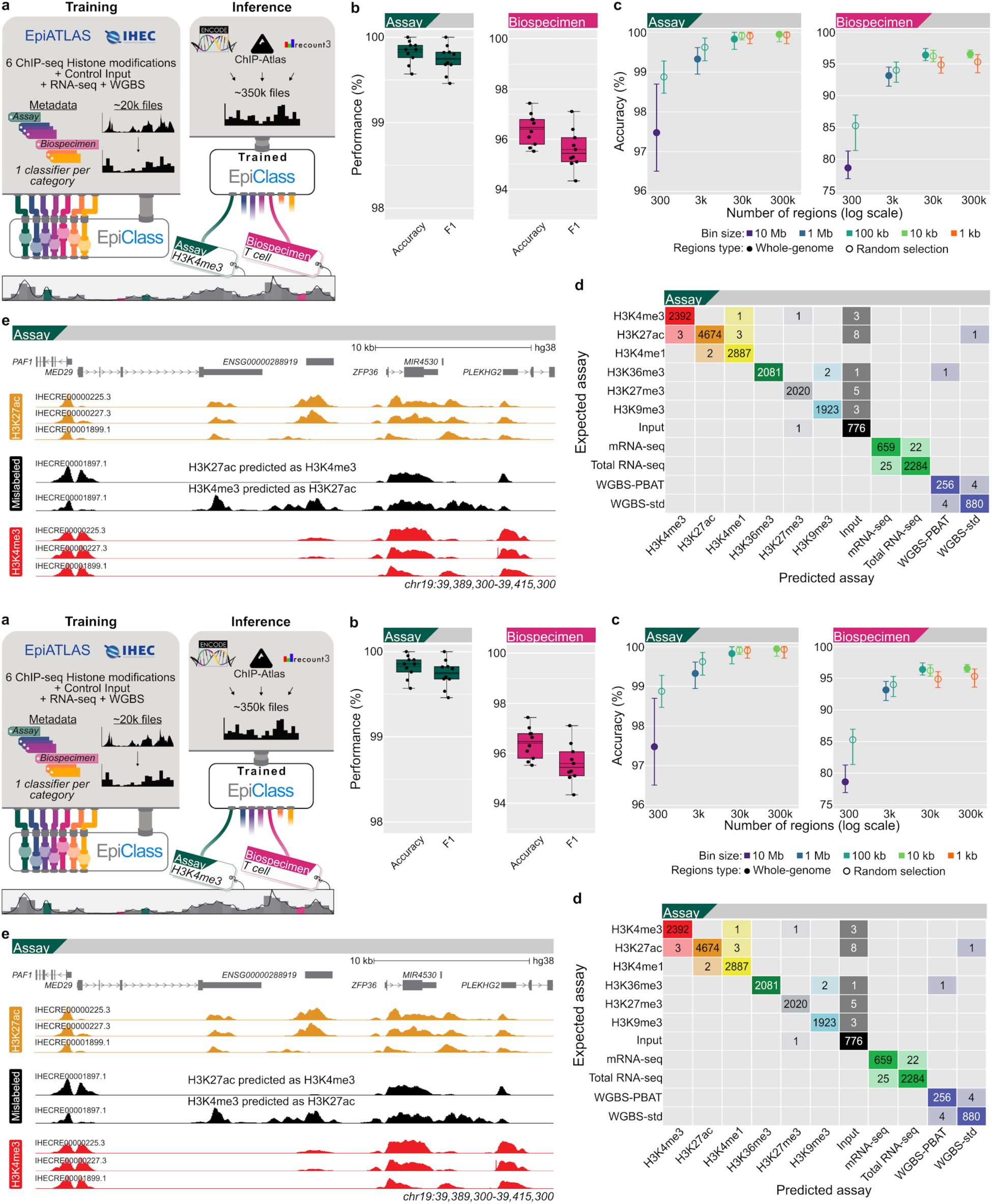
Performance of EpiClass Assay and Biospecimen source classifiers. **a**) Overview of the EpiClass training process for various classifiers and their inference on external data. Each classifier is trained independently. **b**) Distribution of accuracy and F1-score for each of the ten training folds (dots) for the Assay and Biospecimen source MLP classifiers. Dashed lines represent means, solid lines the medians, boxes the quartiles, and whiskers the farthest points within 1.5× the interquartile range. **c**) Distribution of accuracy per training fold for different bin resolutions for the Assay and Biospecimen source classifiers. The circles represent the means and the whiskers the min and max values of the ten training folds (detailed view in Supplementary Fig. 2A-B). Covering the whole genome (filled circles) with non-overlapping bins of 10 Mb requires 315 regions (purple), while it requires 3,044 regions of 1 Mb (blue), 30,321 regions of 100 kb (turquoise), and 303,114 regions of 10 kb (green). To correctly isolate the impact of the bin size, smaller regions were randomly subsampled to match region numbers (open circles), including 1 kb bins (orange). **d**) Confusion matrix aggregating the cross-validation folds (therefore showing all files) without applying a prediction score threshold. RNA-seq and WGBS data were both separated according to two protocols during initial training (but combined thereafter to nine assays). **e**) Genome browser representation showing in black the datasets swap between H3K4me3 and H3K27ac for IHECRE00001897 in the metadata freeze v1.0, along with typical correct datasets over a representative region.

Using five different machine learning approaches, we evaluated classification performance through stratified 10-fold cross-validation. Input features consisted of the average signal computed over 100 kb non-overlapping genome-wide bins, resulting in 30,321 features per signal file (excluding the Y chromosome) (Methods). The Assay classifiers achieved ∼99% accuracy, F1-score, and Area Under the Curve of the Receiver Operating Characteristic (AUROC), while the Biospecimen source classifiers reached ∼95% across the 16 most abundant classes comprising 84% of the epigenomes (the remaining being distributed in 46 smaller classes ignored) (Fig. 1b, Supplementary Fig. 1a-c, Supplementary File 1).

The Multi-Layer Perceptron (MLP, or dense feedforward neural network) showed marginally superior performance on the more complex biospecimen source classification task having important class-imbalance (certain classes being either over or under-represented) (Supplementary Table 2). As our primary goal was to establish a proof-of-concept for this approach, we selected the MLP and focused our subsequent efforts on assessing the model’s performance across different genomic resolutions, rather than on exhaustive hyperparameter optimization. Further analysis with this approach revealed that larger genome-wide bins (1 Mb and 10 Mb) substantially decreased performance, while smaller bins (10 kb and 1 kb) offered minimal improvements despite greatly increasing computational demand (Fig. 1c, Supplementary Table 2). Additional data preprocessing steps, including blacklisted region removal and winsorization, showed no important impact on performance (Supplementary Fig. 1d, Supplementary Table 3, Supplementary File 2, Methods), leading us to adopt the 100 kb bins resolution without further filtering to simplify subsequent analyses.

We also evaluated alternative input feature sets including protein coding genes (∼68 kb on average), cis-regulatory elements showing high correlation between H3K27ac level and gene expression (avg. ∼2.3 kb)^27^, and highly variable DNA methylation segments (200 bp)^28^. For both Assay and Biospecimen source classifiers, none of these alternative feature sets improved the average accuracy by more than 1% compared to the 100 kb bins, with the notable exception of WGBS data. In this case accuracy improved substantially from 85% with 100 kb bins to 93% when using smaller bin sizes and more relevant features (Supplementary Fig. 2, Supplementary Table 2, Supplementary File 3). These findings validated our choice of using the 100 kb approach as an effective compromise, providing comprehensive genome-wide coverage without introducing selection bias from predefined regions, while maintaining strong classification performance and simplifying data processing.

Interestingly, the confusion matrix of the Assay classifier revealed that the very few prediction errors of some individual files occur mainly in specific scenarios: they arise between different protocol types of RNA-seq (mRNA vs total RNA) and WGBS (standard vs PBAT), they involve misclassifications with control Input datasets, or they occur between the activating histone marks (H3K27ac, H3K4me3, H3K4me1) that are typically localized around promoters/enhancers (Fig. 1d). These confusion patterns are all biologically understandable given the functional similarities. For the ChIP and RNA-seq assays, the vast majority of prediction scores exceeded 0.98 and are above 0.9 for biospecimen source prediction, with much lower scores for Input and WGBS assays as expected at the chosen resolution (Supplementary Fig. 1e-f, Supplementary File 4). Importantly, the classifier performances are positively correlated with the prediction scores, which enables the use of these scores as a reliable confidence metric (Supplementary Fig. 1h-i). Increasing the prediction score threshold consistently improves performance, though the scores should not be interpreted as calibrated probabilities.

EpiClass demonstrated practical utility during the development phase by identifying eleven datasets with potentially incorrect assay annotation. After reviewing our findings, data generators examined their original datasets and decided to correct one sample swap between two datasets, and excluded eight contaminated datasets from subsequent EpiATLAS versions (Fig. 1e, Supplementary Fig. 3, Supplementary Table 4, Supplementary Methods). The Assay classifier also validated imputed ChIP datasets from EpiATLAS, achieving perfect predictions and very high prediction scores across all assays (Supplementary Fig. 1g, Supplementary File 5, Methods). Additionally, EpiClass contributed to the identification of 134 low-quality ChIP datasets that were also excluded by the EpiATLAS harmonization working group through notably low prediction scores (or high Input prediction score), indicating noisy signal (Supplementary Fig. 4, Supplementary Table 4).

### EpiClass reliably validates and augments other categories of EpiATLAS metadata

Building on the successful classification of assay and biospecimen source attributes, we extended our approach to six additional categories of harmonized metadata from EpiATLAS (donor sex, sample cancer status, donor life stage, biomaterial type, paired-end sequencing status and data provider (consortium)). These new classifiers demonstrated similarly impressive performance, achieving accuracy above 95%, F1-scores exceeding 92% (except for life stage), and AUROC values greater than 98% (Fig. 2a-b, Supplementary Fig. 5a, Supplementary Table 2). The F1-score variations among categories primarily reflected underlying class imbalances, quantified through normalized Shannon entropy (Fig. 2c, Supplementary Table 1). For instance, in the life stage category, fetal and newborn classes contained only 44 and 106 datasets respectively, compared to over 5,000 adult datasets. Similarly, in the sex category, the mixed class included just 114 datasets versus more than 3,400 for both female and male classes. Despite these class imbalance challenges, the robust classifiers performance enabled us to augment with high-confidence over 85% of previously missing donor sex and life stage labels in EpiATLAS v2.0 metadata, endorsed by the presence of multiple assays per biological sample. Moreover, the performances of all classifiers improve by increasing the prediction score threshold, prompting us to consider prediction scores as a confidence score hereafter (Supplementary Fig. 6a).

**Figure 2.**
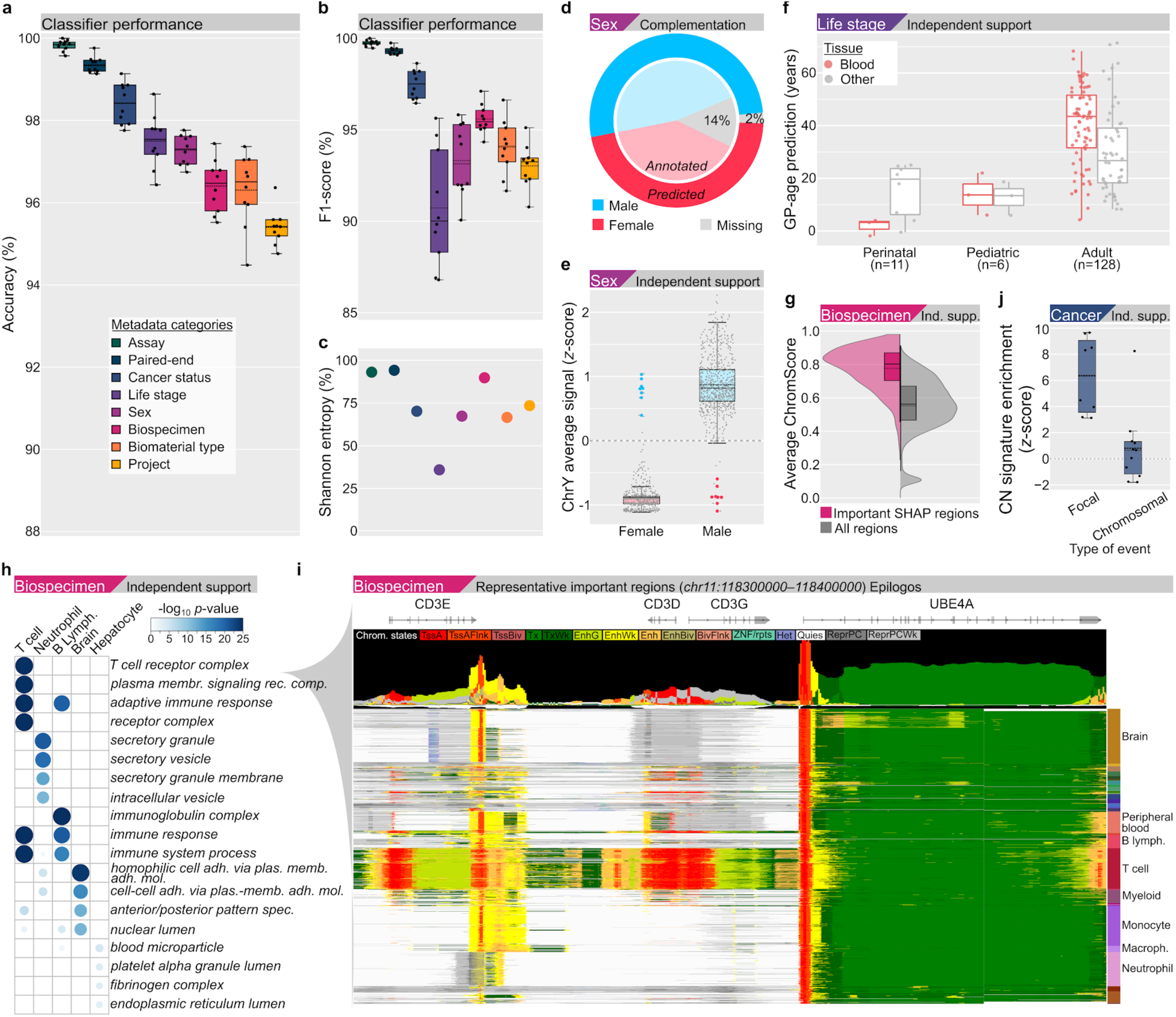
EpiClass reliably validates and augments other categories of EpiATLAS metadata. **a-b**) Distribution of accuracy and F1-score evaluated per training fold (dots) for each metadata classifier. Performance metrics are reported without applying a prediction score threshold. Dashed lines represent means, solid lines the medians, boxes the quartiles, and whiskers the farthest points within 1.5× the interquartile range. **c**) Normalized Shannon entropy scores for each metadata category. **d**) Proportion of donor sex metadata originally annotated (inner circle, metadata v1.1) and predicted with high-confidence (outer circle, metadata v2.0) for female (red) and male (blue) (mixed sex not shown). **e**) Distribution of the average *z*-score signal of epigenomes (dots) over chrY, computed on the ChIP-seq datasets (up to 7 assays per epigenome) using the fold change track type files for female (red) and male (blue). Originally mislabeled epigenomes are shown as big dots. Dashed lines represent means, solid lines the medians, boxes the quartiles, and whiskers the farthest points within 1.5× the interquartile range. **f**) Distribution of the age prediction from GP-age for the WGBS datasets with an originally unknown life stage predicted by EpiClass (datasets related to blood biospecimen source are in red, others in grey). Dashed lines represent means, solid lines the medians, boxes the quartiles, and whiskers the farthest points within 1.5× the interquartile range. **g**) Distribution of the average ChromScore values over the important Biospecimen source classifier regions according to SHAP (pink, n=2,994) compared to the global distribution (grey, n=30,321). Dashed lines represent means, solid lines the medians, boxes the quartiles, and whiskers the farthest points within 1.5× the interquartile range, with a violin representation on top. **h**) Top four gene ontology (GO) terms from the biological process and cellular component categories, enriched for genes within important Biospecimen source classifier regions according to SHAP values by g:Profiler^36^. **i**) Epilogos visualization of one of the important Biospecimen source classifier regions enriched in the T cell receptor complex GO term from panel h. **j**) Distribution of the *z*-score enrichment for copy number (CN) signatures (dots) in genomic regions identified as important by the Cancer status classifier compared to random control regions. CN signatures are grouped as being mostly associated with focal changes (CN9-12,17-21) or chromosomal ones (CN1-8,13-16). Dashed lines represent means, solid lines the medians, boxes the quartiles, and whiskers the farthest points within 1.5× the interquartile range.

To support the sex predictions, we leveraged chromosome Y (chrY) signal as a complementary validation metric given that it was excluded during training. As expected, this signal showed distinct patterns between male and female samples^29^, where males exhibited higher average methylation signals on chrY than females, with the exception of WGBS (Supplementary Fig. 5b). Moreover, progressively increasing the prediction confidence threshold resulted in a corresponding increase in the average *z*-score separation between the clusters, indicating that the prediction score aligns well with the strength of this independent biological sex marker (Supplementary Fig. 5c). Through this approach, we confidently assigned sex metadata to 268 unlabeled biological samples, reducing unknown cases to 2% (Fig. 2d, Supplementary Table 5). This analysis also revealed 23 previously mislabeled samples, which were corrected in EpiATLAS v2.0 (Fig. 2e, Supplementary Table 5, Supplementary Methods).

We achieved similarly strong results for life stage classification, confidently assigning metadata labels to 404 biological samples, with only 5% remaining unknown (Supplementary Table 5). To support these predictions, the WGBS data was provided to the machine learning GP-age tool,^19^ showing expected trends between its predicted age and the life stage categories across both previously unknown (n = 145) and all samples (n = 565) (Fig. 2f, Supplementary Fig. 5d). Notably, blood-related samples showed tighter correspondence between GP-age predictions and life stage categories, with more distinct age distributions across the perinatal, pediatric, and adult groups. This improved alignment is consistent with GP-age’s training background which is made of blood samples. This analysis also identified 17 mislabeled biological samples, which were also corrected in EpiATLAS v2.0 (Supplementary Table 5).

Using Shapley additive explanations (SHAP)^30^, we identified genomic regions that most significantly influence model predictions for biospecimen source classification (Supplementary Methods). SHAP values quantify each feature’s contribution to model decisions by measuring its impact on predictions compared to expected values. This approach revealed an average of 187 important genomic bins driving the Biospecimen source classifier decisions (Supplementary Table 6). Interestingly, their potential biological relevance is supported by two aspects: 1) their elevated functional potential compared to all regions of the genome, quantified using the ChromActivity computational framework^27,31^ (Fig. 2g, Supplementary Fig. 7, Supplementary Methods), and 2) their biospecimen source-specific enriched gene ontology terms that are well aligned with expected ones, also supported by the Epilogos visualization^32^ of a representative region (Fig. 2h-i, Supplementary Table 7, Supplementary Methods).

We also identified 503 SHAP-based important regions for the Sex classifier and 336 for the Cancer status one (Supplementary Table 6). Unsurprisingly, ∼30% of the important regions for sex classifications were on chrX (5.8 fold enrichment), including the known sex-influenced genes XIST^33^ and FIRRE^34^ (*P* = 3.30E-3, Supplementary Methods) (Supplementary Fig. 5e), with X-linked inheritance (HP:0001417) emerging as the most significant term from the 643 protein-coding genes in these regions (*P* = 3.40E-55) (Supplementary Table 7). Notably, 78.6% (22/28) of the features overlapping the pseudoautosomal region 1 (PAR1) were among the important regions, a highly significant enrichment (*P* = 2.55E-18). The important regions from the Cancer status classifier showed a much higher enrichment (*z*-score > 3) in copy number (CN) alteration signatures associated with focal events, such as the tandem duplicator phenotype (CN17), complex patterns (CN18-21) and focal loss of heterozygosity (CN9-12), compared to chromosomal events (Fig. 2j, Supplementary Table 8, Supplementary Methods)^35^. Enriched gene ontology terms from these same important regions are related to chromatin structure (GO:0030527, *P* = 3.38E-23) and histone deacetylation (REAC:R-HSA-3214815, *P* = 2.60E-19) (Supplementary Table 7).

Collectively, these results demonstrate that EpiClass can be used to validate and enrich the metadata of associated data within EpiATLAS, and suggest that the regions identified as important by the different classifiers contain biologically relevant information.

### Application of EpiClass on public epigenomic data

Considering the great performance of EpiClass on EpiATLAS data, we decided to evaluate this approach on external data. We therefore trained six new classifiers on metadata for which equivalent information could be usually extracted from public sources (Assay, Biospecimen source, Sex, Cancer status, Biomaterial type and Life stage) and used the complete EpiATLAS data for training (Methods).

To validate these new classifiers, we first applied them to 4,716 ‘core’ ENCODE datasets generated from the same 9 assays used for training, after excluding biological samples (EpiRR) present in EpiATLAS. A comparison of biospecimen ontology terms revealed that 73% of all ENCODE samples share an exact biospecimen source with EpiATLAS, consistent with the substantial overlap observed between the two resources in PCA space (Supplementary Fig. 8a). As expected, 96% of these datasets have high-confidence scores for the Assay classifier, with an average accuracy and F1-score of ∼98%, while the accuracy ranges from 87% to 95% for the other classifiers with overall ∼90% high-confidence predictions (Fig. 3a, Supplementary Fig. 8b, Supplementary Table 9, Supplementary File 6). Based on these results, a list of 59 potentially contaminated or mislabeled datasets was submitted to the consortium who decided to flag them in the ENCODE Data Portal (Supplementary Table 10). While this does not demonstrate generalization to a large fraction of novel biospecimen sources, it remains a meaningful independent validation as none of the selected individual biological samples overlap between the two resources.

**Figure 3.**
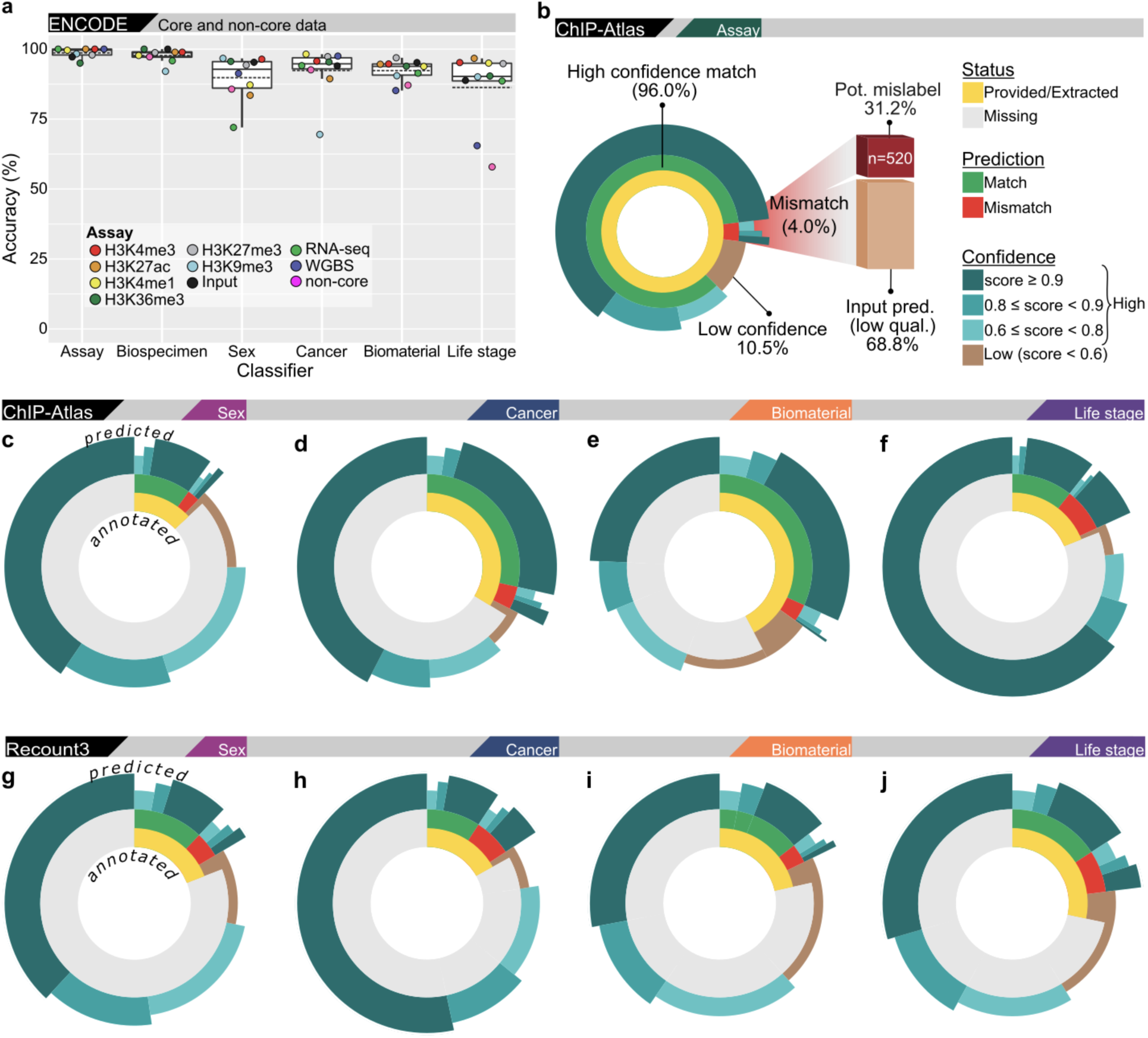
EpiClass results on public datasets. **a**) Accuracy of each metadata classifier breakdown per assay (dots) of the high-confidence predictions on the ENCODE data for which metadata was available, excluding those in EpiATLAS (Supplementary Table 9). Dashed lines represent means, solid lines the medians, boxes the quartiles, and whiskers the farthest points within 1.5× the interquartile range. **b**) Inference of the Assay classifier on the ChIP-Atlas datasets represented as a multi-layer donut chart where the internal layer is depicting the availability status of the metadata as being provided/extracted (yellow) vs missing (grey), the central layer depicting the classifier prediction matching (green) or not (red) the expected metadata label, and the external layer containing the confidence level where prediction scores above 0.6 correspond to high-confidence predictions (light to dark blue >0.6, >0.8 and >0.9) vs low confidence predictions (brown). The high-confidence mismatching predictions were further divided between datasets predicted as Input (light orange) vs potential mislabels (dark red). **c-j**) Inference of the donor Sex (c,g), sample Cancer status (d,h), Biomaterial type (e,i) and donor Life stage (f,j) classifiers on the ChIP-Atlas (c-f) and recount3 (g-j) sources represented as multi-layer donut charts as in panel b.

We next tested the classifier’s robustness on 4,061 ChIP-seq datasets also downloaded from ENCODE but targeting 238 unique ‘non-core’ factors (2,725 datasets) corresponding to proteins and histone modifications different from the six core histone marks. Based on their biological functions, these non-core datasets were manually categorized as being involved in processes happening at different locations relative to genes such as transcriptional regulation, polycomb repression and splicing (Supplementary Table 11). Despite never seeing these specific factors during training, the Assay classifier assigned high prediction scores to ∼75% of these non-core datasets, correctly mapping the vast majority to the functionally related core histone mark (Supplementary Fig. 8c, Supplementary File 6, Supplementary Methods). Unsurprisingly, these non-core factors generally overlap with core datasets in a Principal Component Analysis (PCA) (Supplementary Fig. 8a). As anticipated based on these observations, the performances of the other classifiers on the non-core datasets are overall comparable to core ones (Fig. 3a, Supplementary Fig. 8d). And very reassuringly, the classifier performances on both types of ENCODE datasets are positively correlated with the prediction scores, increasing the confidence towards all these classifiers (Supplementary Fig. 6b-c). Interestingly, the EpiClass prediction score of the Input class for non-Input datasets is a good indicator of sample quality based on ENCODE ChIP-seq quality metrics, for both core and non-core datasets (Supplementary Fig. 8e-f, Supplementary Table 12, Supplementary Methods). Overall, these results demonstrate the generalization ability of the Assay classifier, recognizing fundamental chromatin patterns and associating unseen ChIP non-core factors with the appropriate functional category represented by the core marks.

We next applied these classifiers to human ChIP-seq datasets from experiments targeting the seven core ChIP assays (six histone modifications plus Input), uniformly reprocessed by ChIP-Atlas that extracted its data and metadata from GEO. However, due to lack of standardization in GEO metadata and the fact that >5,100 entries were not labeled in ChIP-Atlas, we cross-referenced these metadata with three other sources: NGS-QC, Cistrome DB, and our independent reinterpretation of GEO’s metadata (Methods). We thus downloaded 47,065 datasets, excluding those identified as non-core assays by any of the sources. It was reassuring that 98% of datasets received identical target annotations across multiple metadata sources (Supplementary Fig. 9a, Supplementary File 7). To avoid confounding performance results, we excluded 1,047 datasets from ChIP-Atlas that were also included in EpiATLAS through ENCODE.

Interestingly, ∼90% of the Assay classifier predictions of the remaining 46,018 datasets were high-confidence, and 96% of them were matching the expected assay (Fig. 3b). Moreover, the Assay classifier was also able to resolve 99% of the 619 high-confidence predictions where the metadata sources disagreed (Supplementary File 7). These results are even more impressive considering that the vast majority of the reprocessed data from ChIP-Atlas do not overlap with EpiATLAS and ENCODE in a PCA (likely because different reprocessing pipelines were used) (Supplementary Fig. 9b).

These predictions also unveiled discrepancies in many datasets, with 520 mislabeled candidates partially validated by manual inspection (Fig. 3b, Supplementary File 7). These potential mislabeled datasets are coming from 224 different GEO series (GSEs) but are not evenly distributed, with 50% of them concentrated in 41 GSEs (18%), including 28 datasets that were apparently swapped by the authors within 8 GSEs (e.g., six H3K27ac registered as H3K9me3 and vice versa in GSE106660) (Supplementary Fig. 9c, Supplementary File 7). More than 2/3 of high-confidence mismatches were predicted as Input, corresponding to low signal-to-noise ratio as demonstrated on ENCODE datasets (Supplementary Fig. 8e-f) and therefore identified as low quality datasets (Fig. 3b). Additionally, to assess the impact of using imputed data, we trained two new assay classifiers restricted to the six core histone marks (no Input) on EpiATLAS: one on observed and one on imputed data. Each classifier was applied to the other EpiATLAS data type and to ChIP-Atlas (Supplementary Fig. 9d, Supplementary File 8). For five of the six marks, both classifiers reached similar per-assay accuracies ranging between 89% and 99%, with differences of at most 1% within EpiATLAS and 4.7% on ChIP-Atlas. H3K27ac was the exception: on ChIP-Atlas, the imputed-trained classifier reached 71.8% accuracy, compared with 86.9% for the observed-trained classifier. This may however partly reflect the under-representation of H3K27ac in the imputed training set (1,088 files vs about 1,700 for each other mark; Supplementary Table 13).

Beyond validating assay labels, we next assessed the classifiers’ ability to augment other key but optional metadata attributes that original authors rarely submit to GEO. Considering the excellent performances of EpiClass classifiers on EpiATLAS and ENCODE datasets, as well as the precision of the Assay classifier on ChIP-Atlas datasets, it was reassuring to observe that 82% of the high-confidence predictions of the Sex classifier match the provided or extracted label, 88% for Cancer status, and 91% for Biomaterial type (Fig. 3c-e, Supplementary Table 9). We were therefore able to augment with high-confidence between 77% and 93% of the corresponding missing metadata. However, the Life stage classifier showed markedly different behavior: only 58% of high-confidence predictions matched the expected label (Fig. 3F). Moreover, increasing the prediction score threshold for the Life stage classifier did not improve performance, unlike the other classifiers (Supplementary Fig. 6D). This indicates that the Life stage classifier is poorly calibrated for this data, prompting us to not recommend new life stage predictions for the ChIP-Atlas dataset. Additionally, the Biospecimen source classifier was not applied to public data, as its training content was not representative of the sample diversity found in these repositories.

Finally, we evaluated EpiClass performance on 316,228 human RNA-seq datasets from recount3 using a new Assay classifier trained this time using all of the nine types of assay from EpiATLAS. As expected, virtually all (99.7%) datasets were predicted as RNA-seq or mRNA-seq with high-confidence (Supplementary Table 9, Supplementary File 9), even though they are showing only partial overlap in the PCA with EpiATLAS and ENCODE (Supplementary Fig. 9e). However, the proportion of these filtered (Methods) high-confidence predictions matching the expected label was only 73% for the Sex classifier, 59% for Cancer status and 82% for Biomaterial type (Fig. 3g-i). Although these matching rates are much lower than the ones obtained on ChIP-Atlas, they remain substantial considering that the vast majority of EpiATLAS data used for the training are stranded files (meaning there is one file per strand per dataset), while the files from recount3 are unstranded (combining the signal from both strands) and include reads mapping at multiple locations; this is affecting the predictions (Supplementary Fig. 5g, Supplementary File 10, Supplementary Methods) and should therefore be taken into account in future training to generate more robust classifiers. The Life stage classifier showed a better accuracy (69%) here than for ChIP-Atlas, increasing this time with the prediction score threshold (Fig. 3j, Supplementary Fig. 6e). Nevertheless, these predictions, as well as those from the Cancer classifier, should be interpreted with more caution considering their overall profile. Still, focusing the analysis specifically on the 6,109 filtered datasets from recount3 reported as leukemia (a cancer type well-represented in the training data), the high-confidence accuracy rose from 59% to 76%, highlighting the impact of the training data (Supplementary Table 9). Selecting the classifiers with >80% label concordance for high-confidence predictions, we were able to augment over 320,000 metadata attributes for the donor sex, sample cancer status and biomaterial type, representing on average 85% of the public datasets for which these information were missing (Supplementary Table 9, Supplementary File 9).

## Discussion

In this study, we introduced EpiClass, a suite of machine learning classifiers designed to address the pervasive challenge of incomplete and inaccurate metadata in public epigenomic repositories. We have demonstrated, as a proof-of-concept, that by leveraging the largest high-quality harmonized data and metadata from the IHEC EpiATLAS project, it is possible to predict fundamental metadata attributes directly from the genomic signal itself. During its development, EpiClass not only proved to be valuable by refining its own training data through the identification of a handful of mislabeled assays (including some released in 2013 by the NIH Roadmap Epigenomics consortium) and low-quality datasets, but also successfully augmented EpiATLAS by enabling the completion of missing metadata for hundreds of biological samples with trustworthy predictions (Fig. 1 and 2). Some of these predictions were supported by independent analyses, including those from the Sex classifier through the use of chrY signal (absent from the original training). The intriguing inverse relationship observed between sex for WGBS datasets compared to the other assays (Supplementary Fig. 5) can be explained by the fact that DNA methylation is typically higher for the pseudoautosomal region 1 (PAR1) than the rest of the X chromosome, in addition to the lower methylation levels of chrY due to its relative transcriptional inactivity and high heterochromatin content^29^. Considering that most reads mapping to chrY in females inevitably originate from the highly methylated pseudoautosomal regions, the WGBS signal is therefore higher than in males where it represents an average of both PAR1 and the rest of chrY. Independent analyses supporting the biological validity of the Life stage and Biospecimen source classifiers predictions also contributed to the relevance of EpiClass. Overall, this work provides the first systematic, signal-based approach for the validation and augmentation of metadata across diverse epigenomic assays.

The generalizability of EpiClass was further validated on unseen ENCODE datasets (Fig. 3a). Notably, the Assay classifier helped identify additional contaminated or mislabeled datasets flagged accordingly in the ENCODE Data Portal (Supplementary Table 10), and also successfully mapped previously unobserved non-core ChIP-seq targets to their most functionally related core histone marks (Supplementary Fig. 8c). This demonstrates that the model has successfully learned fundamental genomic localization patterns driven by shared underlying chromatin biology, rather than dataset-specific features This is consistent with epigenomic data being known to cluster strongly by both assay and biospecimen source^27,37^.

The potential utility of EpiClass was further explored through its application to over 350,000 public ChIP-seq and RNA-seq datasets (Fig. 3b-j). This large-scale analysis validated the reported assay for the majority of ChIP-Atlas datasets while also flagging hundreds of potential mislabels, such as dataset swaps and low-quality experiments. Beyond validation and correction, this effort highlighted the potential for metadata augmentation, particularly for fundamental but often omitted information like donor sex and sample cancer status. However, we observed that prediction accuracy varied considerably depending on the classifier and the data source (Supplementary Fig. 6). Using filtering based on an 80% average accuracy threshold applied per data source and per classifier we could complement metadata for a substantial portion of previously unlabeled datasets (between 47% and 97%), contributing more than 320,000 key metadata attributes, primarily from donor sex and biomaterial type. The performance of some of the other classifiers proved more fragile when applied to more diverse public datasets, indicating that their utility is currently best suited for data more closely matching the training distribution. This challenge is well exemplified by the Life Stage classifier. Its accuracy dropped significantly on public datasets most probably because its training was dominated by a vast overrepresentation of adult datasets. This severe class imbalance rendered the model poorly calibrated for the more diverse life stage distributions found in public repositories. Conversely, the performance of the Cancer status classifier on the leukemia datasets from recount3 demonstrates the positive impact of data similarity. These examples highlight a critical principle for the application of EpiClass and machine learning in general: a model’s performance is intrinsically linked to the composition of its training data. Consequently, while promising, significant work remains to develop classifiers that are robust across the full spectrum of public data (e.g., adding multi-mapped reads for RNA-seq as for recount3, diversifying the biospecimen sources), underscoring that these predictions must be interpreted with an understanding of the model’s limitations. Classifiers trained on imputed data reached accuracies similar to those trained on observed data for most histone marks, suggesting that imputed datasets could help expand training sets for future assay classifiers (Supplementary Fig. 9d). However, whether imputation preserves the sample-level features needed for attributes such as the donor sex or biospecimen source remains to be evaluated.

Our methodological choices were key to the success reported above, but they also come with considerations. The use of 100 kb non-overlapping genomic bins covering the whole genome proved to be a pragmatic and effective compromise, capturing sufficient information for accurate classification without the prohibitive computational cost and therefore environmental carbon footprint of smaller resolutions. This bin size is also biologically relevant considering the average protein-coding gene length of 68 kb (Methods), and it was also used in other studies^29^. It is also interesting to note that the performance of the 100 kb bins were similar to the 1 MB bins, which is similar to the size of Topologically Associating Domains (TADs) that range from a few hundred kilobases to a couple of megabases, with a mean size of around 650 kb in humans^38^.

While our SHAP analysis successfully identified biologically relevant genomic regions driving predictions, such as biospecimen source-relevant GO enrichment, the XIST locus identification for sex classification and focal CN-associated regions for cancer status (Fig. 2g-j and Supplementary Fig. 5e), we acknowledge that SHAP can have limitations in the presence of highly correlated features^39^, which are ubiquitous in genomics. Future work could explore alternative feature attribution methods like the Marginal Contribution Feature Importance^40^ to further refine our understanding of the models’ decision-making processes.

Ultimately, this work serves as a powerful proof to the value of large-scale data harmonization efforts. The existence of EpiATLAS, with its standardized processing and meticulous metadata curation^41^, was the essential foundation upon which EpiClass was built. Our classifiers not only enhance the quality and reliability of existing public data for key metadata types, but also pave the way for a more complete and accurate future for data repositories. The next logical step is to train new, more robust models by leveraging the predictions from this study passing stringent quality thresholds (often known as ‘silver standard’ vs ‘gold standard’). Furthermore, it would be relevant to extend to a wider array of ChIP-seq targets and other assays like ATAC-seq and single-cell data, in addition to supporting other model organisms and developing user-friendly interfaces. Such a tool would ideally be integrated into data submission pipelines, providing real-time feedback to researchers and allowing them to correct potential errors at the source, therefore contributing to the long-standing challenge recognized by the GEO developers regarding the metadata standardization^1^. By enriching the metadata of public epigenomic datasets, EpiClass directly supports data reusability, a core aim of the FAIR principles (findable, accessible, interoperable, and reusable data), and helps unlock the potential of public genomic data for biological discovery. Our work not only enhances the quality and reliability of epigenomic data, but also underscores the value of harmonized metadata in ensuring the integrity and enriching large-scale public datasets.

## Methods

### Input data and preprocessing for EpiClass training

This study utilizes 20,922 epigenomic BigWig signal files from 7,464 independent experiments (datasets) conducted on 2,216 biological samples of the EpiATLAS project, corresponding to the files included in the metadata freeze v1.1 of EpiATLAS with some minor differences as detailed in Supplementary Table 14 and 15. Moreover, 9,570 ChIP-seq imputed BigWig signal files from EpiATLAS were also used for some training (Supplementary Fig. 9d) and inference (Supplementary Fig. 1g) tasks (Supplementary Table 13, Supplementary File 5). The average signal over given bins was computed on each signal file and converted to the Hierarchical Data Format version 5 (HDF5) format using the epiGeEC tool^37^. Each chromosome, excluding chrY for historical reasons, was processed separately then all concatenated. The bin at the end of a chromosome is typically smaller than the other bins. The signal in each file is centered and scaled globally (as *z*-score) prior to classifier training or inference.

A key metadata field, (’track_type’, specifies the nature of the signal data within each HDF5 file, which varies depending on the assay:

#### ChIP-seq assays

typically three track types are available per dataset: raw read counts, fold change enrichment over corresponding control Input, and the statistical significance of enrichment expressed as a p-value rejecting the null hypothesis that the signal at that location is present in the control. Corresponding Input control datasets are by definition represented only by the raw read counts (’ctl_raw’) track type.

#### RNA-seq assays

the data primarily consists of strand-specific raw read counts (’Unique_plusRaw’ for the positive strand, ‘Unique_minusRaw’ for the negative strand), with a smaller number of merged-strand raw count tracks (’Unique_raw’) also present. The training was conducted by providing the two protocol types of RNA-seq (mRNA vs total RNA) independently, but the results were often merged to simplify the results.

#### WGBS assays

strand-specific signals are provided as ‘gembs_pos’ and ‘gembs_neg’ tracks. The training was conducted by providing the two protocol types of WGBS (standard vs Post-Bisulfite Adaptor Tagging (PBAT)) independently, but the results were often merged to simplify the results.

These different track types provide signal files representing variations of the same underlying dataset (experiment) and are treated accordingly in downstream steps like cross-validation fold creation and oversampling (see Training and evaluation strategy). Multiple tracks of the same dataset possess the same Universally Unique Identifier (UUID).

### Machine learning model architecture and selection

Our classification analysis employed five distinct machine learning approaches, each chosen for specific capabilities:

A Multi-Layer Perceptron (MLP) Neural Network formed our primary model, featuring a single fully connected hidden layer of 3,000 nodes. This architecture implements ReLU activation functions and employs both dropout and weight decay (using the Adam Optimizer) for effective regularization, a technique that helps models generalize better by discouraging overfitting to dataset-specific features or noise.

For comparison and validation, we implemented four additional classifiers: a Random Forest (RF), a Logistic Regression (LR) with softmax output function, a Support Vector Machine (SVM) with linear kernel, and a gradient boosting machine (GBM). While we considered alternative SVM kernels such as RBF and polynomial, memory constraints limited us to the linear kernel implementation.

For the RF, LR and SVM, scikit-learn^42^ was used. For the GBM, we selected the LightGBM implementation^43^ using the DART booster method^44^ for its built-in regularization properties, which reduced the need for extensive hyperparameter optimization.

### Hyperparameter Optimization

The complete set of hyperparameters used during the 10-fold cross-validation performance evaluation is provided in Supplementary Table 16, and a summary of the hyperparameter tuning procedure is given in Supplementary Table 17.

A distinct strategy was applied to MLP versus non-neural network models. For the RF, LR, SVM, and LightGBM classifiers, hyperparameters were tuned on a 90% subset of the full training set, split into nine folds. The hyperparameter configuration yielding the best average validation performance (across one held-out fold at a time) was then used for the final 10-fold cross-validation evaluation on the complete training data. For RF, LR, and SVM, hyperparameters were optimized using Bayesian optimization (BayesSearchCV from *scikit-optimize*), with 30 iterations per fold (i.e., 30 candidate hyperparameter sets tested per fold). For LightGBM, a different strategy was used: key hyperparameters (feature_fraction, num_leaves, bagging_fraction) were tuned sequentially with the Optuna^45^ (stepwise tuning, i.e. optimizing one parameter at a time in sequence, following the Optuna documentation, for a total of 12 parameter sets). This procedure ensured that all baseline models operated near their optimal configurations for comparison.

For the MLP, we deliberately refrained from large-scale, systematic hyperparameter optimization. Initial exploratory tests varied the learning rate, dropout rate, weight decay strength, batch size, and the number/size of hidden layers. These tests showed that:

1. The chosen architecture (a single hidden layer of 3000 nodes) provided sufficient capacity, with larger architectures offering little benefit or even hindering convergence.
2. Learning rate and batch size primarily influenced convergence speed rather than final accuracy.
3. The selected dropout and weight decay settings yielded effective regularization without suppressing performance.

Crucially, the MLP achieved competitive or superior performance compared with the tuned classifiers even with this fixed configuration. Given the high computational cost of systematic neural network tuning and the study’s aim to benchmark the MLP as a strong baseline, we focused on this architecture for direct comparison against the tuned models. This approach highlights the robustness and effectiveness of the selected MLP configuration for the classification task.

During cross-validation, training was allowed to proceed for up to 300 epochs, with early stopping applied if validation accuracy failed to improve for 60 epochs. Regularization (dropout and weight decay) effectively prevented overfitting, even when the validation performance plateaued for hundreds of epochs. The final MLP models used for inference on external datasets were trained for the full 300 epochs on the entire EpiATLAS dataset, because no validation set was available to enable early stopping.

### Training and evaluation strategy

We evaluated model performance using a stratified 10-fold cross-validation strategy designed to prevent data leakage. Our approach leverages the structure of the EpiATLAS data, which is organized by the Epigenome Reference Registry (EpiRR). An EpiRR (https://www.ebi.ac.uk/epirr) groups all datasets derived from a given biological source, ensuring they share consistent metadata such as donor sex, cancer status and life stage.

To prevent data leakage, we constrained our cross-validation splits at the UUID level, assigning all track types from a single dataset to the same fold. This ensures that the model cannot learn to recognize a particular dataset from partial information, for example by seeing one signal track (e.g., fold change) during training and predicting on another track (e.g., p-value) derived from the same assay. However, because UUIDs are nested within EpiRRs, different datasets generated from the same biological source may appear in separate folds. This leaves open the possibility of residual information leakage between related assays, such as histone modifications that tend to co-localize, where shared epigenomic patterns from the same biological source could provide the model with indirect signal, or allow it to recognize the sample itself, which would inflate the prediction of sample-level labels. Constraining the splits at the EpiRR level would have removed this risk, but retraining every classifier and rederiving every SHAP related result would have been computationally costly. As a sanity check that the UUID-level grouping did not introduce meaningful leakage and affected model performances, we retrained two classifiers with an EpiRR-level split (otherwise trained identically), in which all datasets from a given epigenome are assigned to the same fold, and compared them with the UUID-level split.

For the Assay and Biospecimen source classifiers at 100 kb resolution, we compared the per-fold metrics (accuracy, macro F1, micro and macro AUC) of the two splitting strategies with two-sided Welch’s t-tests (n = 10 folds per strategy, 8 uncorrected tests). Both are exposed to leakage, through different mechanisms: assay labels differ between the datasets of an epigenome, but correlated assays from the same sample share epigenomic patterns, whereas biospecimen source labels, like other sample-level labels, are shared by all datasets of an epigenome. The other categories were not retrained with EpiRR-level grouping. Tests designed for cross-validated comparisons, such as the 5×2cv paired t-test^46^ and the corrected resampled t-test^47^, compare classifiers on shared train/test partitions; they do not apply here because the two conditions differ by their partition. The folds of a cross-validation share most of their training data, so their metrics are positively correlated, and Welch’s test likely underestimates the variance of the difference. This makes the test anti-conservative, biased toward detecting differences, so the absence of significant differences is not an artifact of the test. None of the comparisons reached significance (all p > 0.05, Supplementary Table 18, Supplementary File 11), suggesting that UUID-level grouping does not introduce meaningful leakage for these categories.

Class imbalance was addressed by randomly oversampling minority classes (with replacement) to match the majority class size within each training fold. Oversampling was performed at the UUID level to preserve related measurements, using a fixed random seed (N = 42) to ensure identical cross-validation folds across classifiers with the same training data.

### Compute requirements

Training time varied depending on the classification task, genomic resolution, number of input features, and the extent of class imbalance, which determined the amount of oversampling required within each training fold. As a representative benchmark, training the assay classifier (11 classes) at 100 kb resolution for 300 epochs required approximately 1 hour on a single NVIDIA A100-SXM4-40GB GPU using native 16-bit Automatic Mixed Precision (AMP). For tasks with more input features and/or stronger class imbalance, training time could increase to approximately 6 hours. These values reflect GPU training time only and exclude data loading and preprocessing overhead, which was typically <10% of total wall time during training. Task-specific runtimes are provided in Supplementary Table 19.

EpiClass is lightweight at inference time and does not require a GPU. A trained 100 kb-resolution MLP model has a checkpoint size of approximately 1.07 GB, which sets the minimum memory requirement for inference. As a benchmark, classifying 10,000 samples (1.38 GB of input data) required approximately 1 minute on a consumer desktop CPU (Intel Core i7-6700, 2015), with approximately 75% of the wall time spent on file loading and preprocessing rather than model execution.

### Task-specific filtering and evaluation

#### Biospecimen source classification

This classifier uses the ‘Intermediate Biospecimen Label’ field (also called ‘harmonized_sample_ontology_intermediate’) in EpiATLAS. During training, biospecimen sources were filtered to retain only labels that include datasets across all EpiATLAS assays and at least 10 unique datasets per biospecimen source-assay combination to ensure at least one dataset per fold; a total of 16 biospecimen sources of the 62 unique labels fulfill that condition.

#### recount3 inference

To ensure that EpiClass metadata classifiers were applied to datasets similar to the training dataset, a filtering step was applied by selecting datasets classified as RNA-seq/ mRNA-seq by the Assay classifier (11 classes) with prediction scores >0.6, therefore excluding ∼78k/316k (25%) datasets; this strategy provided more usable data than only using the ∼151k directly labeled as RNA-seq datasets (48% of total).

#### Life stage classification

Datasets with a biomaterial type of cell-line were excluded as they lack meaningful developmental classifications and were absent from training data. Although trained on five labels, three early categories (embryonic, fetal, newborn) were merged into ‘perinatal’ for external evaluation, as the highly imbalanced training dataset provided insufficient resolution to distinguish between these stages on dissimilar external data. Indeed, F1-scores for these classes were very low even on ENCODE data, the closest dataset to EpiATLAS Since ChIP-Atlas and recount3 had few provided non-cell-line datasets labeled (∼9% and ∼1% respectively), we applied our biomaterial classifier to increase the number of datasets on which the Life stage classifier was applied. We used prediction score thresholds that achieved ≥90% recall for explicitly labeled cell-line datasets (0.6084 for ChIP-Atlas, 0.8465 for recount3). Notably, since these filtered datasets constitute the majority, performance metrics are more representative of the complete set compared to using only explicitly labeled non-cell-line datasets. We did not apply this filtering to datasets already possessing explicit biomaterial labels. The additional datasets are likely closer to the training distribution (in-distribution datasets, i.e., datasets with characteristics similar to training data) due to this filtering approach.

### Performance metrics

Model performance for each classification task was assessed using several standard metrics computed on the validation sets created within the cross-validation (one metric value per fold). The scikit-learn (python) implementations of the following metrics were used:

#### Accuracy

The overall proportion of correctly classified files or datasets (per fold, per class or globally, sometimes also breakdown per assay where only the files of a given assay are used).

#### F1-score

The unweighted average of the F1-scores calculated per class (also called macro F1-score). The F1-score, defined as the harmonic mean of precision and recall, provides a balance between these two metrics, and the macro-average ensures equal contribution from each class, making it less sensitive to class imbalance than accuracy.

#### Area Under the ROC Curve (AUROC)

Assessed using the One-Versus-Rest (OvR) scheme. We report both micro-AUROC (computed globally by considering all elements of the confusion matrix) and macro-AUROC (unweighted average of per-class AUROCs). The macro-AUROC under the OvR scheme can still be sensitive to class imbalance due to the changing composition of the ‘rest’ group for each class.

Prediction scores produced by the classifiers (except for SVMs, which only output the predicted class) can serve as an approximate measure of prediction confidence. These scores can be aggregated by UUID or by EpiRR to obtain a single consensus value (Supplementary Fig. 1e,f). In a ChIP-seq dataset for example, each UUID has three associated files (one per track type), while in a given EpiRR with nine assays there could be a total of 23 associated files (18 for the six ChIP-seq, one for the corresponding Input, and two each for RNA-seq and WGBS).

There are two ways to derive a consensus prediction from the class and score of each file:

- Majority vote first: identify the most frequently predicted class among the files, then average the prediction scores for only those files.
- Highest average score first: calculate the average prediction score for each predicted class and select the class with the highest mean, regardless of how many files were predicted for it.

When averaged prediction scores are reported, the first method (majority vote followed by averaging) is used.

### The effect of outlier values

To initially assess the impact of potential outlier features on model performance, a series of input data transformations were applied sequentially (Supplementary Fig. 1d). Supplementary File 2 details each prediction.

#### Blacklist Filtering

At 1 kb resolution, all feature positions overlapping with blacklisted genomic regions^48^ were set to zero. This step removes data from regions prone to sequence alignment artifacts, which may bias model learning. Following this, feature values were averaged as usual to obtain data at 100 kb resolution.

#### Winsorization

On the blacklist-filtered 100 kb resolution data, winsorization was applied within each file to the top 0.1% of feature values across the genome. These values were clipped to the 99.9th percentile, thereby limiting the influence of extreme outliers that may otherwise cause overfitting. The effects of winsorization alone, or of applying winsorization prior to blacklist filtering, were not evaluated in this study.

### Alternative model input features

#### Genomic resolution

To assess the effect of resolution on model performance, genomic bins were subdivided or merged by factors of 10, yielding resolutions from 10 Mb to 1 kb (Fig. 1c). For resolutions from 10 Mb to 10 kb, the entire genome was covered uniformly, varying only the bin size and feature count. At 1 kb resolution, where ∼3 million features exceeded the GPU memory available at this time (32 GB), random subsampling was used to match the 303,114 regions of the 10 kb baseline. This enabled approximate performance comparisons while controlling for feature set size.

#### Feature selection

Other biologically relevant sets of regions were selected to evaluate their impact on training (Supplementary Fig. 2). The gene set contains 19,864 regions corresponding to protein-coding genes with 2 kb flanking regions, derived from gencode.v29 annotations^49^ (Supplementary File 12), with an average gene length of approximately 68 kb.

Two other sets of ∼30k and ∼300k features use the cis-regulatory elements (obtained from ChromHMM) showing high correlation between H3K27ac level and neighbor gene(s) expression from the EpiATLAS flagship paper^27^. The regions selected are those with the highest maximum absolute Spearman correlations (Supplementary File 13).

Two other sets of ∼30k and ∼300k features represent highly variable methylation segments. The mean was calculated across all CpGs within each of the 15,155,223 bins of 200 bp provided by EpiATLAS for all datasets (EpiATLAS data, DNA_Methylation_segmentation/epilogos/ scores_*.txt.gz). If a dataset has at least one missing (NA) value for a CpG, its mean value was set to NA. The variance was then calculated on the means of the 9,906,813 bins containing <5% of NA. The segment type assignment of each bin is described elsewhere^28^. As expected, about half of the ∼30k and ∼300k selected segments were from the Lowly Methylated Regions (LMR) type that is mostly biospecimen source-specific, corresponding to an enrichment of ∼18x compared to overall segments distribution. Due to a technical error, only the signal from the first 2 bp of the 200 bp from the most variable DNA methylation segments were actually used to generate the HDF5 files (Supplementary File 14), but the overall correlation with the desired signal over the full 200 bp bins was still high (∼0.8), and the resulting classifier performances (with 2 bp) were highly similar to using the most variable CpG 2 bp regions (not shown).

The harmonized EpiATLAS data, including all files previously mentioned in this subsection, is available to download here: https://ihec-epigenomes.org/epiatlas/data/.

### ChIP-seq metadata extraction from four public databases

The metadata information of human ChIP-seq datasets from the following four public databases was all downloaded/updated in July 2023.

#### Chip-Atlas

a single source file of metadata was downloaded (https://chip-atlas.dbcls.jp/data/metadata/experimentList.tab) and parsed with an internal script to extract relevant information (https://github.com/GFrosi/ChIP-Atlas-extraction).

#### Cistrome DB v2.0

a series of json files were downloaded (http://cistrome.org/db) and parsed (https://github.com/GFrosi/CistromeDB_Metadata).

#### NGS-QC

a series of html files were extracted (https://ngsqc.org/database.php) and parsed (https://github.com/GFrosi/NGS-QC-extraction.com/GFrosi/NGS-QC-extraction).

#### GEO

the NCBI’s E-utilities^50^ was used to programmatically extract the metadata in XML files and parsed with an internal script to extract relevant information (https://github.com/GFrosi/XML-Parse-GEO-NCBI). The SRX identifiers were used to link the ChIP-Atlas metadata to GSM and GSE identifiers used in the other databases and the target/assay was further standardized (https://github.com/GFrosi/Comparing_Databases_GEO). To circumvent the absence of standardization in GEO metadata, we applied a dictionary of regular expressions (including different names and antibodies catalog numbers of the histone modifications and the control Input) to the ChIP-antibody, Target, GSM title and Source cell fields.

### ChIP-Atlas data download

After cross-referencing the metadata of ChIP-seq data from the four previous sources, we downloaded 50,493 BigWig files from ChIP-Atlas in July 2023 as described in their instructions (https://github.com/inutano/chip-atlas/wiki#downloads_doc). A total of 47,065 ChIP-seq and control Input were successfully converted into HDF5 using the epiGeEC tool^37^ and further used in this study. This number excludes datasets identified as non-core assays by any of the sources.

### ENCODE metadata extraction and data download

The metadata was extracted/updated in February 2025 using the ENCODE REST API (https://www.encodeproject.org/help/rest-api/). The file accessions were linked to the experiment, biosample, biosample type, and file metadata. The following metadata categories were used as is for evaluating classification performance: donor sex (BIOSAMPLE_sex) and donor life stage (BIOSAMPLE_life_stage). The assay was extracted from FILE_target and EXPERIMENT_assay_title. The biospecimen source ontology was linked to EpiATLAS v’harmonized_intermediate_sample_ontology’ via BIOSAMPLE_TYPE_term_id. The biomaterial type (BIOSAMPLE_TYPE_classification) had slight remapping (’tissue’ → ‘primary tissue’) and the following terms were ignored: ‘in vitro differentiated cells’ and ‘organoid’. The cancer status was extracted from 15 fields, using a dictionary of cancer related terms and their associated class (https://github.com/labjacquespe/EpiClass/tree/v0.8.3/src/R/ENCODE).

The ENCODE data was downloaded in March 2023 from the official portal (https://www.encodeproject.org/), either as BigWig format for the core and non-core datasets, or as BAM for the corresponding control ChIP Input files. The core data correspond to the six histone modifications used by EpiATLAS (p-value track type), RNA-seq (total and polyA, ‘plusUniq’ and ‘minusUniq’ track types) and WGBS (plus and minus track types), while the non-core mainly correspond to transcription factors and histone modifications different from the six core. The BAM files from control Input were converted to BigWig using the IHEC containerized ChIP-seq reprocessing pipeline (https://github.com/IHEC/integrative_analysis_chip/blob/dev-organize-output/encode-wrapper/postprocess/postprocess.sh). A total of 1,782 ChIP-seq, 1,284 control Input, 1,552 RNA-seq and 98 WGBS as well as 4,061 non-core files were successfully converted into HDF5 using the epiGeEC tool^37^ and further used in this study.

### recount3 metadata extraction and data download

Metadata from human datasets identified from the ‘available_samples’ function was retrieved using the recount3 R package in December 2024^51^. The ‘sra.sample_attributes’ column was used to extract metadata for every category. The assay was derived using the additional fields ‘recount_pred.curated.type’, ‘recount_pred.pattern.predict.type’, and ‘recount_pred.pred.type’, while information on biomaterial type, cancer status, and biospecimen source ontology was extracted using the additional field ‘sra.sample_title’.

The project and sample ID were used to generate URLs to download the BigWig files of every dataset from the ‘sra’ source. A total of 316,228 RNA-seq datasets were successfully converted into HDF5 using the epiGeEC tool^37^ and further used in this study.

## Supporting information

Supp. Material Inventory

## Data availability

The harmonized EpiATLAS data is available from the portal (https://epigenomesportal.ca/ihec/), and the metadata from GitHub (https://github.com/IHEC/epiATLAS-metadata-harmonization/ blob/v1.1.1/openrefine/v1.1/IHEC_metadata_harmonization.v1.1.extended.csv). Reprocessed public datasets were downloaded from their respective data portal.

## Code availability

All code for model training and figure reproduction is available on GitHub (https://github.com/labjacquespe/EpiClass/tree/v0.10.5/). For figure reproduction python notebooks, see “fig[X].ipynb” files in the “paper-final” folder. Supplementary Table 19 provides training details for each classifier, including the specific code commit used and links to the corresponding Comet.ml training logs. Cross-validation predictions can be downloaded from the linked Comet.ml pages under ‘Assets & Artifacts/others’. Models used for predictions on ENCODE, ChIP-Atlas, and recount3 databases (see "Application of EpiClass on public epigenomic data") are available on Hugging Face under the EpiClass models collection (https://huggingface.co/collections/KatLeChat/epiclass-models). For the latest version of interactive figures and associated code, visit the Quarto website: https://labjacquespe.github.io/EpiClass/epiclass-paper/index.html

## Acknowledgements

We thank: IHEC for the harmonized epigenomic data and metadata access and Integrative Analysis Working Group for helpful discussions; Marcel H. Schulz and Sébastien Rodrigue for critical reading of the manuscript; Zakaria Méliane for help downloading some datasets; the *Centre de Calcul Scientifique de l’Université de Sherbrooke*, *Calcul Québec* and the Digital Research Alliance of Canada for providing support and access to advanced research computing resources. This work was supported by the Natural Sciences and Engineering Research Council (NSERC) of Canada to P.-É.J. [#435710]. J.R. is the recipient of a Graduate Scholarship from NSERC and the *Fonds de Recherche du Québec – Nature et Technologie* (FRQNT). P.-É.J. holds a *Fonds de Recherche du Québec – Santé* (FRQS) Research Scholar Senior Career Award.

## Author contributions

P.-É.J. and J.L. conceptualized the study; P.-É.J. and J.R. designed the study; J.R. developed the majority of the EpiClass software (data management, classifier implementation, and training/ cross-validation pipeline), and analyzed results; G.F. performed initial download, preprocessing and metadata management, and participated in results analysis; F.W. conducted in-depth metadata processing, analyzed data, and prepared most figures based on results and drafts from J.R.; J.L. prototyped the EpiClass training pipeline; P.-É.J. supervised the project, analyzed and interpreted results, and wrote the manuscript with J.R.

## Inventory of Supporting Information

- Supplementary Methods;
- 10 Supplementary Figures;
- 21 Supplementary Tables;
- 14 Supplementary Files.

An explicit inventory of the supplementary tables and supplementary files is available as an xlsx file.

---

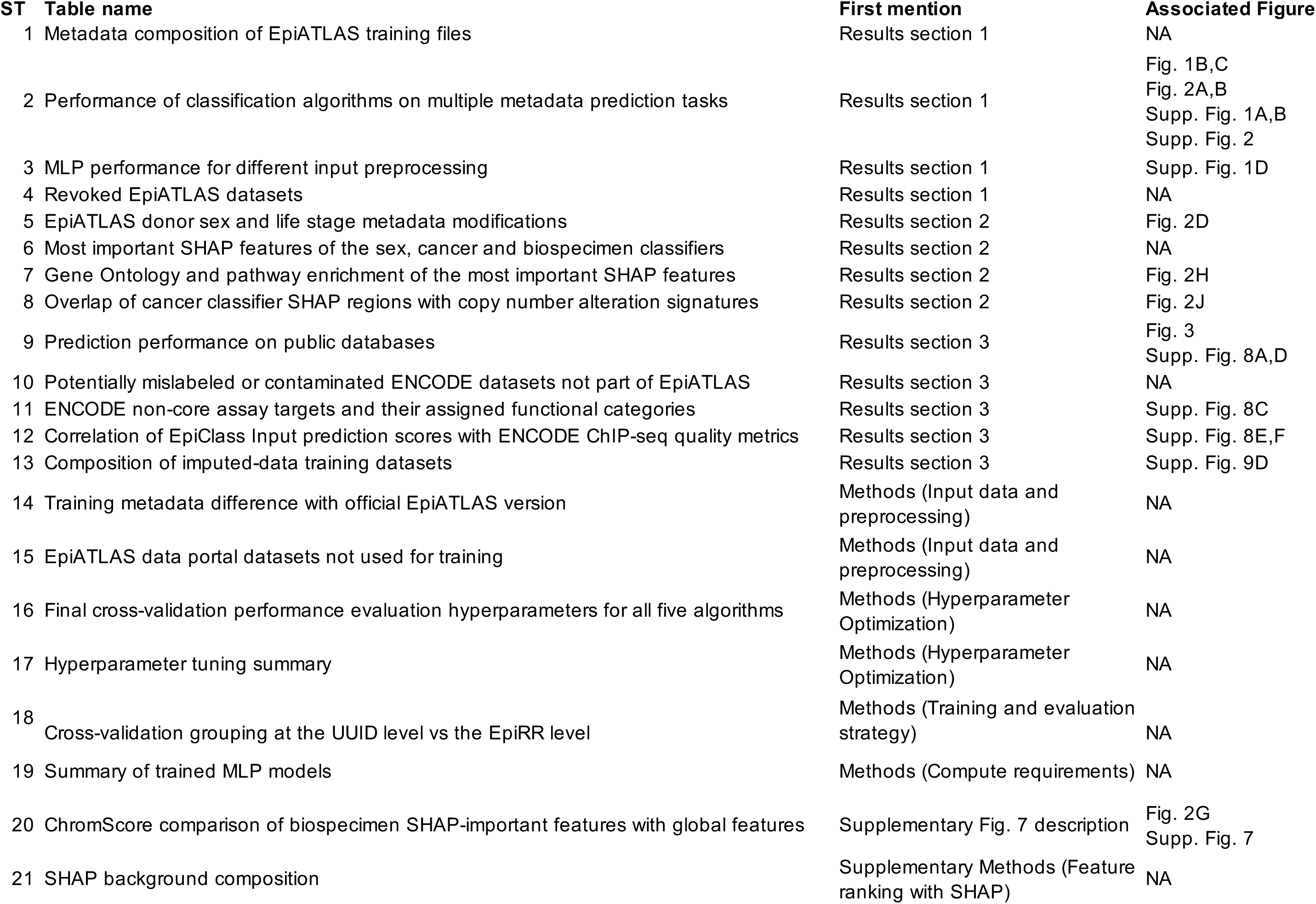

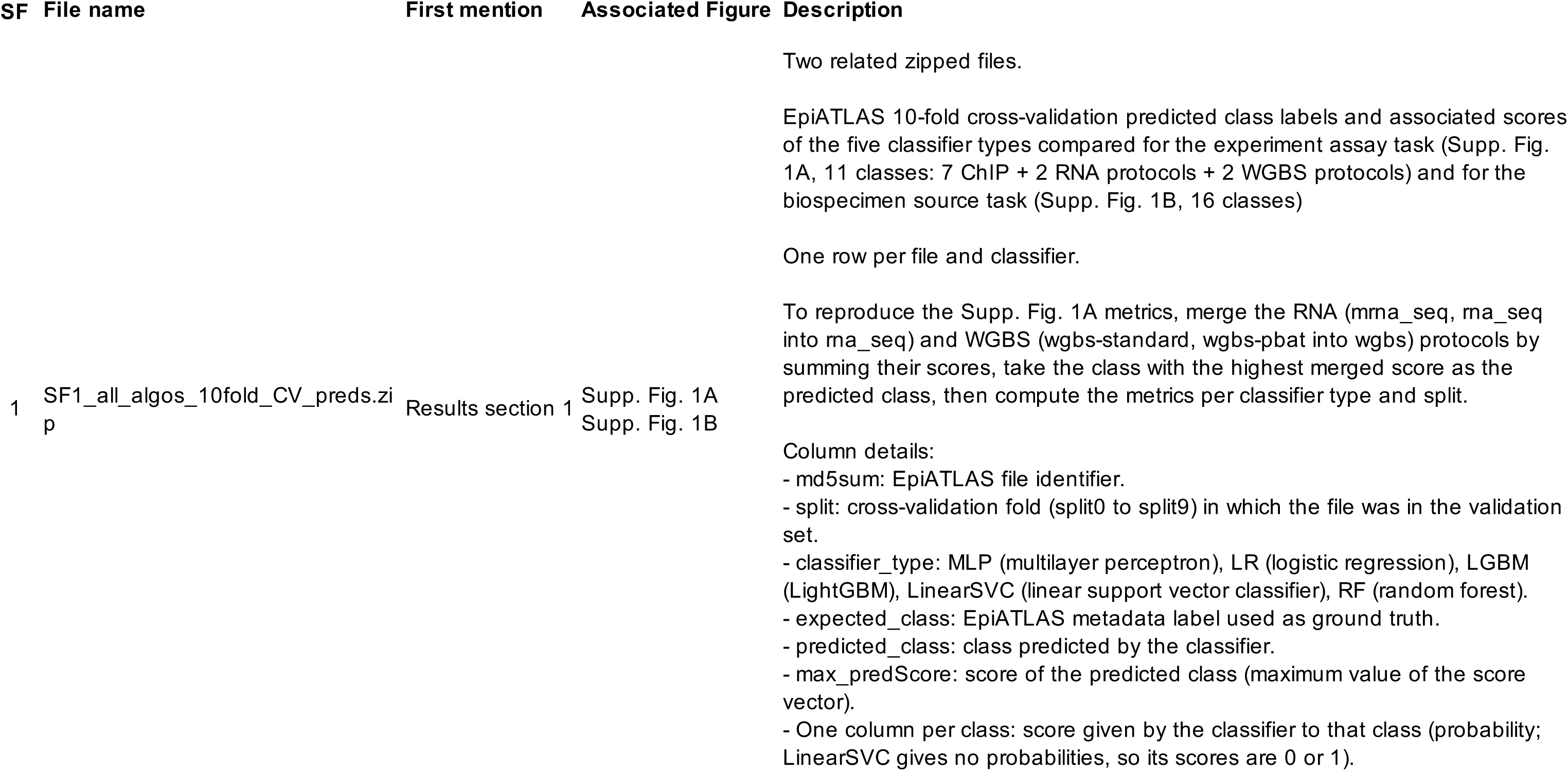
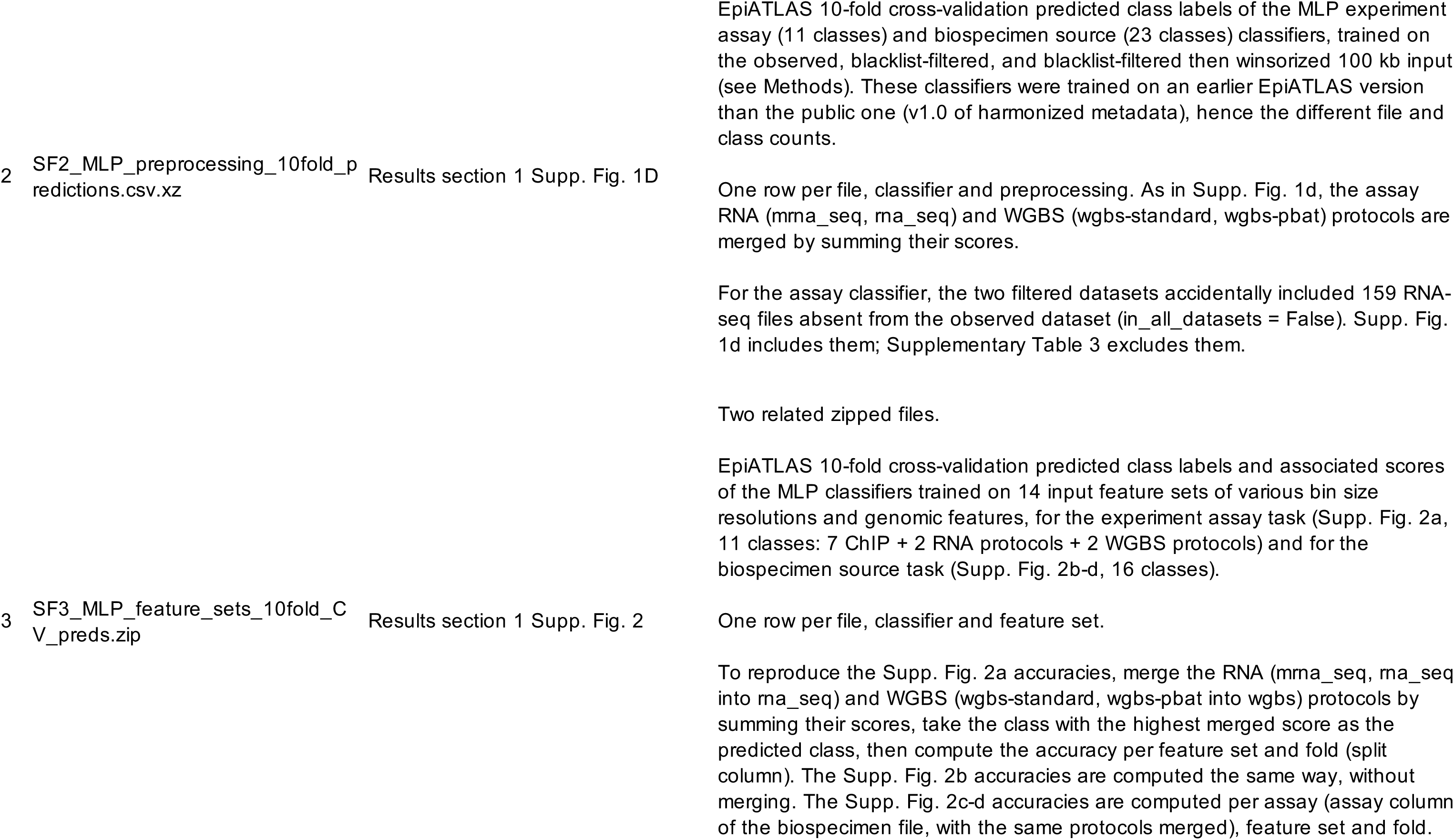
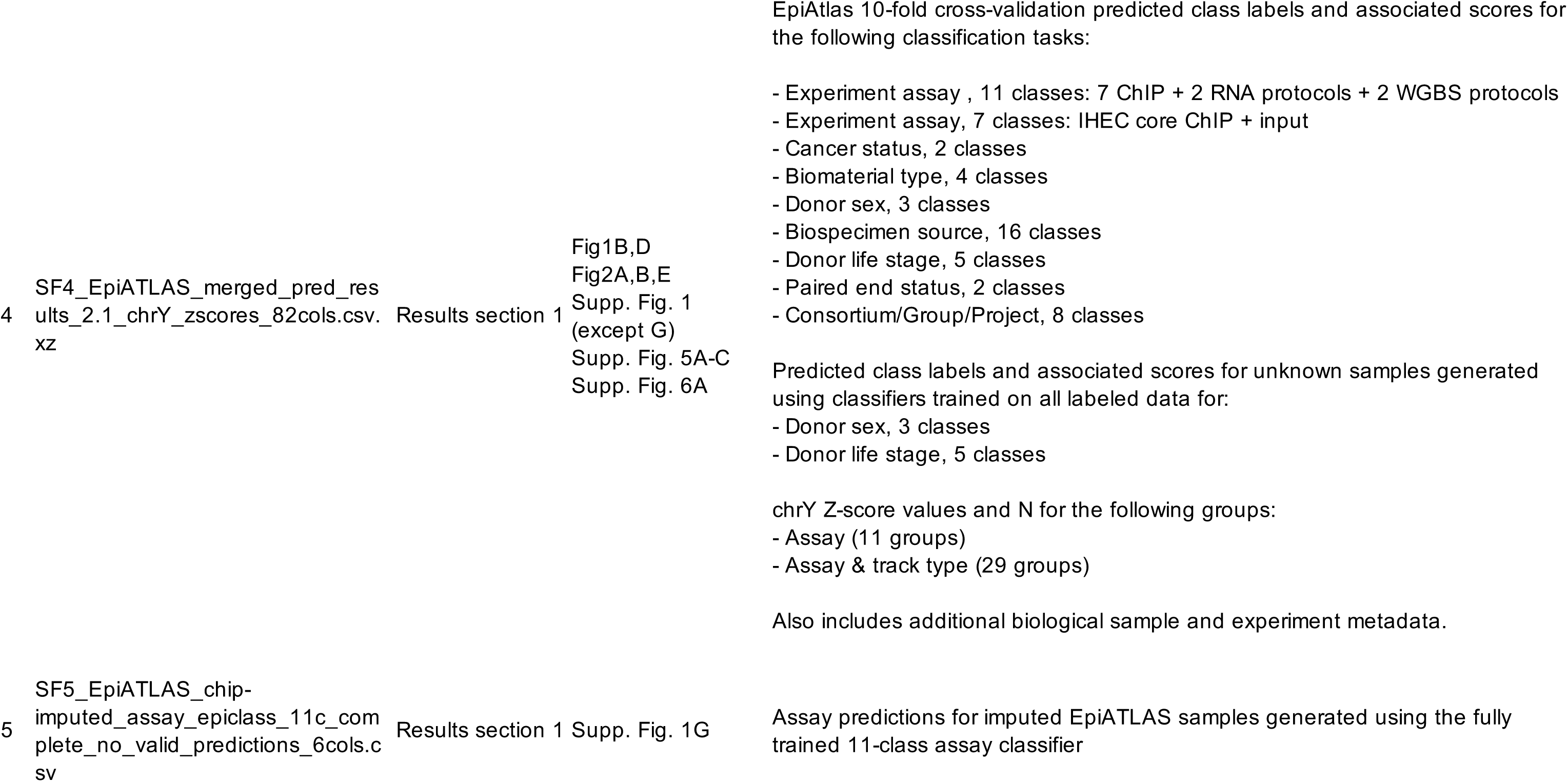
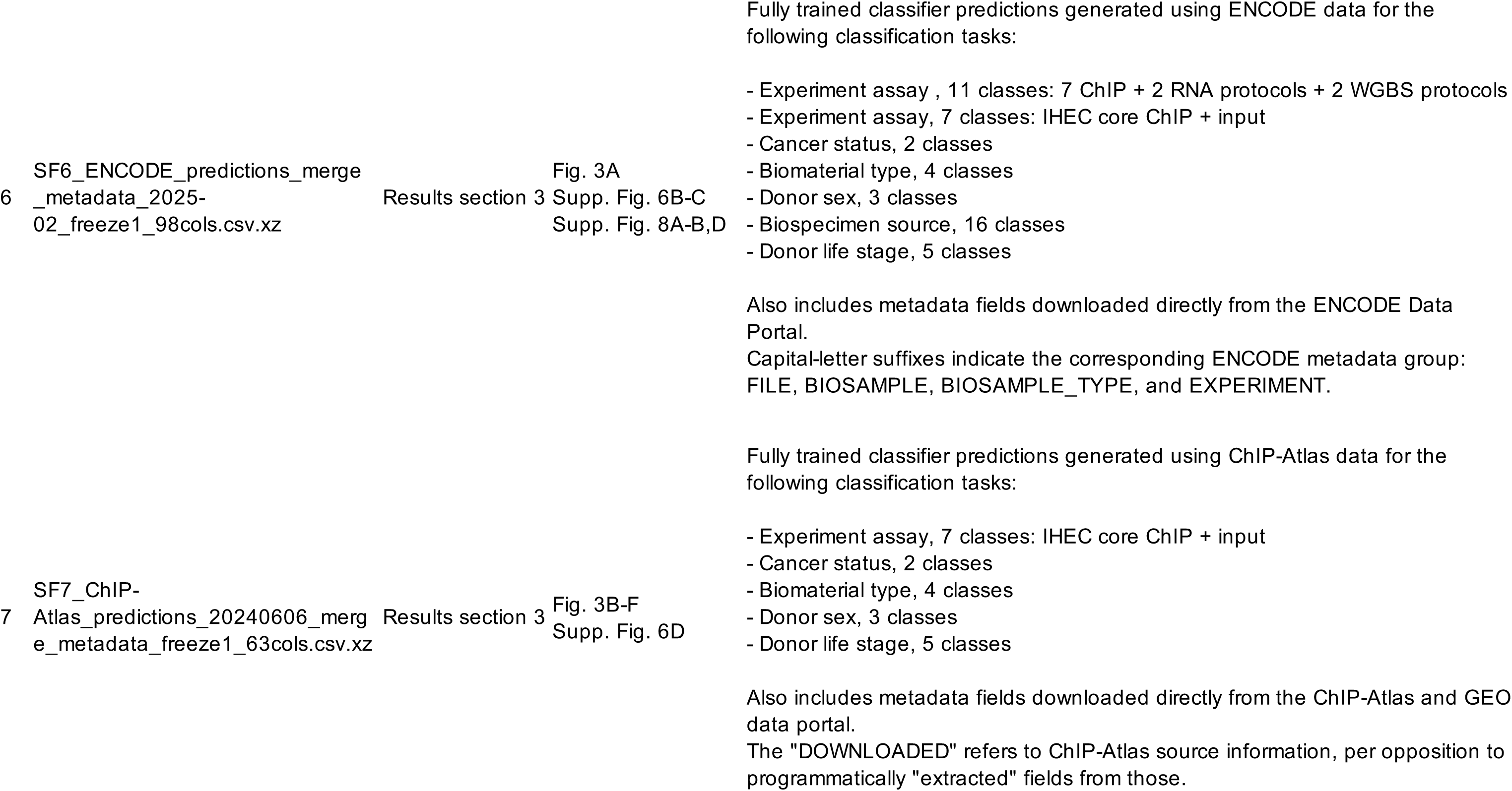
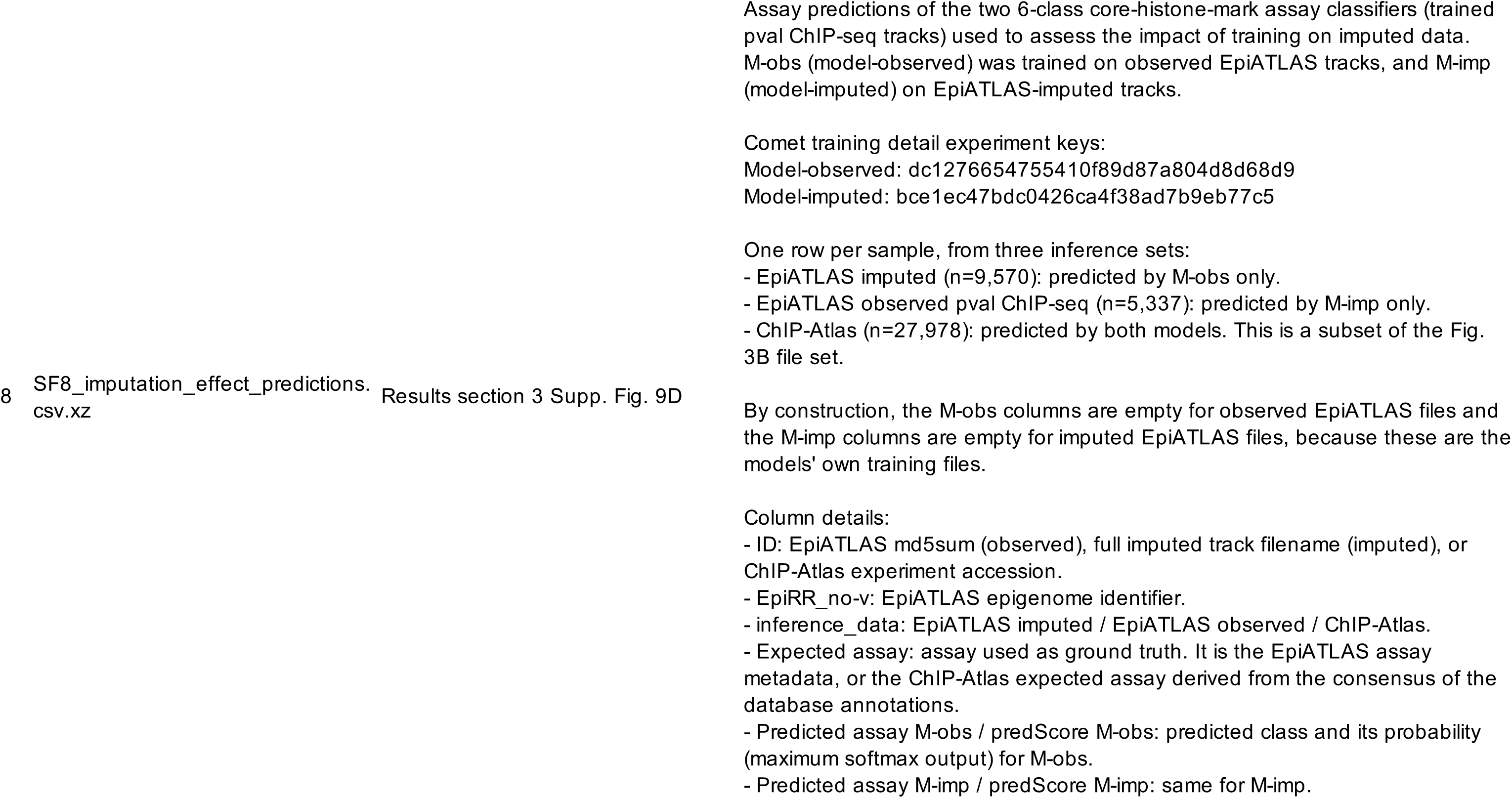
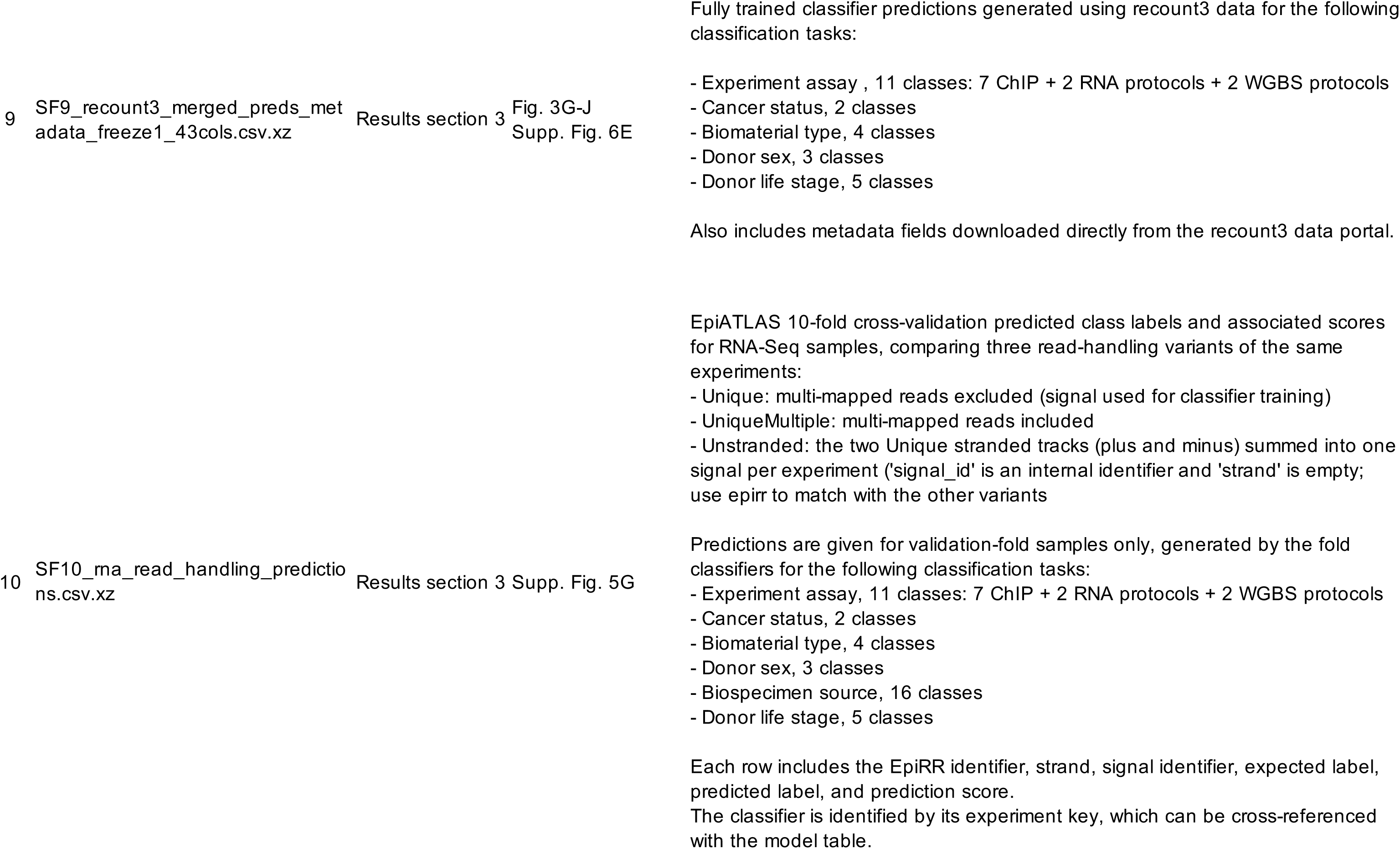
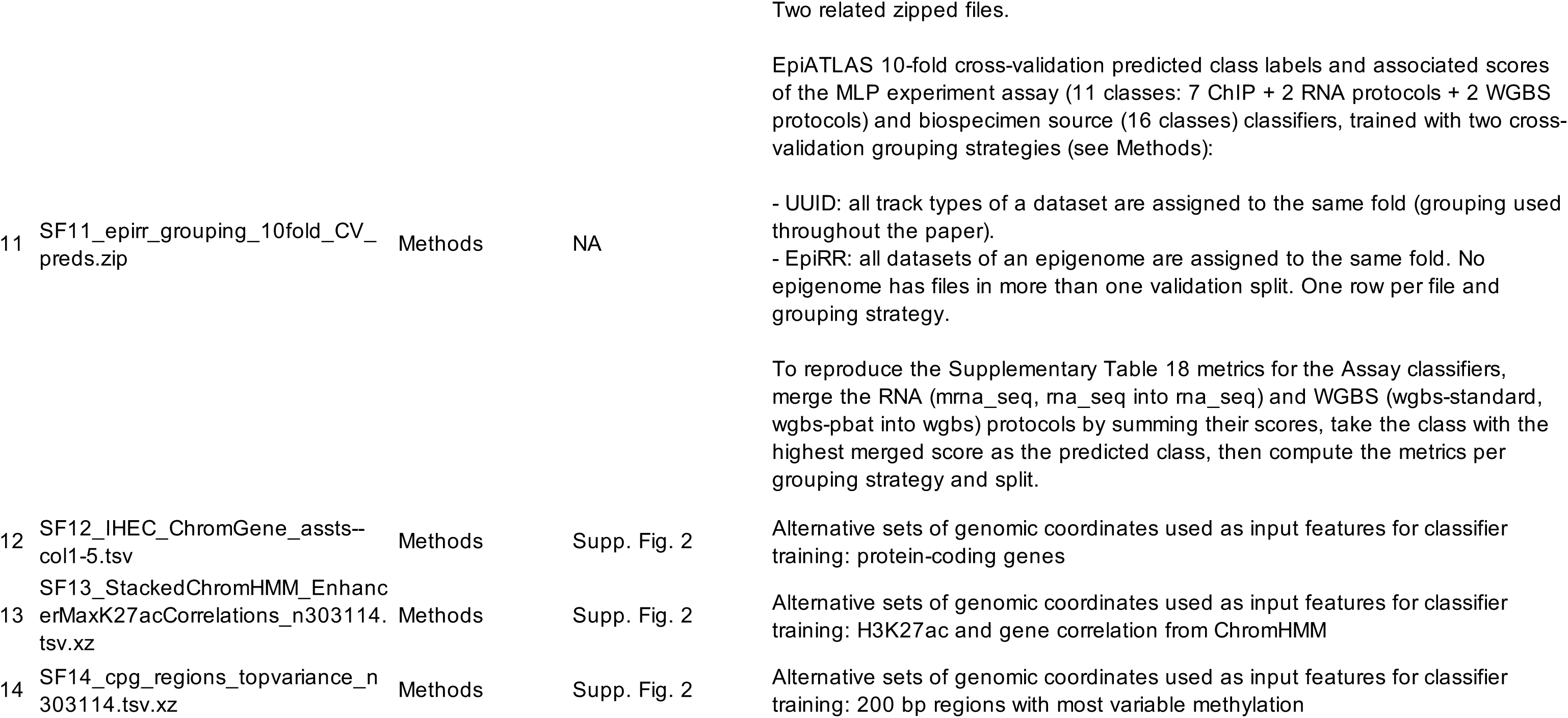

