## Supplementary material for "Leveraging a large harmonized epigenomic data collection for metadata prediction to validate and augment over 350,000 public epigenomic datasets": Supp. Material Inventory

| ST | Table name | First mention | Associated Figure |
|---|---|---|---|
| 1 | Metadata composition of EpiATLAS training files | Results section 1 | NA |
| 2 | Performance of classification algorithms on multiple metadata prediction tasks | Results section 1 | Fig. 1B,C<br>Fig. 2A,B<br>Supp. Fig. 1A,B<br>Supp. Fig. 2 |
| 3 | MLP performance for different input preprocessing | Results section 1 | Supp. Fig. 1D |
| 4 | Revoked EpiATLAS datasets | Results section 1 | NA |
| 5 | EpiATLAS donor sex and life stage metadata modifications | Results section 2 | Fig. 2D |
| 6 | Most important SHAP features of the sex, cancer and biospecimen classifiers | Results section 2 | NA |
| 7 | Gene Ontology and pathway enrichment of the most important SHAP features | Results section 2 | Fig. 2H |
| 8 | Overlap of cancer classifier SHAP regions with copy number alteration signatures | Results section 2 | Fig. 2J |
| 9 | Prediction performance on public databases | Results section 3 | Fig. 3<br>Supp. Fig. 8A,D |
| 10 | Potentially mislabeled or contaminated ENCODE datasets not part of EpiATLAS | Results section 3 | NA |
| 11 | ENCODE non-core assay targets and their assigned functional categories | Results section 3 | Supp. Fig. 8C |
| 12 | Correlation of EpiClass Input prediction scores with ENCODE ChIP-seq quality metrics | Results section 3 | Supp. Fig. 8E,F |
| 13 | Composition of imputed-data training datasets | Results section 3 | Supp. Fig. 9D |
| 14 | Training metadata difference with official EpiATLAS version | Methods (Input data and preprocessing) | NA |
| 15 | EpiATLAS data portal datasets not used for training | Methods (Input data and preprocessing) | NA |
| 16 | Final cross-validation performance evaluation hyperparameters for all five algorithms | Methods (Hyperparameter Optimization) | NA |
| 17 | Hyperparameter tuning summary | Methods (Hyperparameter Optimization) | NA |
| 18 | Cross-validation grouping at the UUID level vs the EpiRR level | Methods (Training and evaluation strategy) | NA |
| 19 | Summary of trained MLP models | Methods (Compute requirements) | NA |
| 20 | ChromScore comparison of biospecimen SHAP-important features with global features | Supplementary Fig. 7 description | Fig. 2G<br>Supp. Fig. 7 |
| 21 | SHAP background composition | Supplementary Methods (Feature ranking with SHAP) | NA |

| SF | File name | First mention | Associated Figure | Description |
| --- | --- | --- | --- | --- |
| 1 | SF1_all_algos_10fold_CV_preds.zip | Results section 1 | Supp. Fig. 1A<br>Supp. Fig. 1B | Two related zipped files.<br><br>EpiATLAS 10-fold cross-validation predicted class labels and associated scores of the five classifier types compared for the experiment assay task (Supp. Fig. 1A, 11 classes: 7 ChIP + 2 RNA protocols + 2 WGBS protocols) and for the biospecimen source task (Supp. Fig. 1B, 16 classes)<br><br>One row per file and classifier.<br><br>To reproduce the Supp. Fig. 1A metrics, merge the RNA (mrna_seq, rna_seq into rna_seq) and WGBS (wgbs-standard, wgbs-pbat into wgbs) protocols by summing their scores, take the class with the highest merged score as the predicted class, then compute the metrics per classifier type and split.<br><br>Column details:<br>- md5sum: EpiATLAS file identifier.<br>- split: cross-validation fold (split0 to split9) in which the file was in the validation set.<br>- classifier_type: MLP (multilayer perceptron), LR (logistic regression), LGBM (LightGBM), LinearSVC (linear support vector classifier), RF (random forest).<br>- expected_class: EpiATLAS metadata label used as ground truth.<br>- predicted_class: class predicted by the classifier.<br>- max_predScore: score of the predicted class (maximum value of the score vector).<br>- One column per class: score given by the classifier to that class (probability; LinearSVC gives no probabilities, so its scores are 0 or 1). |

| | | | |
|---|---|---|---|
| 2 | SF2_MLP_preprocessing_10fold_predictions.csv.xz | Results section 1 Supp. Fig. 1D | EpiATLAS 10-fold cross-validation predicted class labels of the MLP experiment assay (11 classes) and biospecimen source (23 classes) classifiers, trained on the observed, blacklist-filtered, and blacklist-filtered then winsorized 100 kb input (see Methods). These classifiers were trained on an earlier EpiATLAS version than the public one (v1.0 of harmonized metadata), hence the different file and class counts.<br><br>One row per file, classifier and preprocessing. As in Supp. Fig. 1d, the assay RNA (mrna_seq, rna_seq) and WGBS (wgbs-standard, wgbs-pbat) protocols are merged by summing their scores.<br><br>For the assay classifier, the two filtered datasets accidentally included 159 RNA-seq files absent from the observed dataset (in_all_datasets = False). Supp. Fig. 1d includes them; Supplementary Table 3 excludes them. |
| 3 | SF3_MLP_feature_sets_10fold_CV_preds.zip | Results section 1 Supp. Fig. 2 | Two related zipped files.<br><br>EpiATLAS 10-fold cross-validation predicted class labels and associated scores of the MLP classifiers trained on 14 input feature sets of various bin size resolutions and genomic features, for the experiment assay task (Supp. Fig. 2a, 11 classes: 7 ChIP + 2 RNA protocols + 2 WGBS protocols) and for the biospecimen source task (Supp. Fig. 2b-d, 16 classes).<br><br>One row per file, classifier and feature set.<br><br>To reproduce the Supp. Fig. 2a accuracies, merge the RNA (mrna_seq, rna_seq into rna_seq) and WGBS (wgbs-standard, wgbs-pbat into wgbs) protocols by summing their scores, take the class with the highest merged score as the predicted class, then compute the accuracy per feature set and fold (split column). The Supp. Fig. 2b accuracies are computed the same way, without merging. The Supp. Fig. 2c-d accuracies are computed per assay (assay column of the biospecimen file, with the same protocols merged), feature set and fold. |

| | | | | |
|---|---|---|---|---|
| 4 | SF4_EpiATLAS_merged_pred_results_2.1_chrY_zscores_82cols.csv.xz | Results section 1 | Fig1B,D<br>Fig2A,B,E<br>Supp. Fig. 1 (except G)<br>Supp. Fig. 5A-C<br>Supp. Fig. 6A | EpiAtlas 10-fold cross-validation predicted class labels and associated scores for the following classification tasks:<br><br>- Experiment assay , 11 classes: 7 ChIP + 2 RNA protocols + 2 WGBS protocols<br>- Experiment assay, 7 classes: IHEC core ChIP + input<br>- Cancer status, 2 classes<br>- Biomaterial type, 4 classes<br>- Donor sex, 3 classes<br>- Biospecimen source, 16 classes<br>- Donor life stage, 5 classes<br>- Paired end status, 2 classes<br>- Consortium/Group/Project, 8 classes<br><br>Predicted class labels and associated scores for unknown samples generated using classifiers trained on all labeled data for:<br>- Donor sex, 3 classes<br>- Donor life stage, 5 classes<br><br>chrY Z-score values and N for the following groups:<br>- Assay (11 groups)<br>- Assay & track type (29 groups)<br><br>Also includes additional biological sample and experiment metadata. |
| 5 | SF5_EpiATLAS_chip-imputed_assay_epiclass_11c_complete_no_valid_predictions_6cols.csv | Results section 1 | Supp. Fig. 1G | Assay predictions for imputed EpiATLAS samples generated using the fully trained 11-class assay classifier |

| | | | | |
|---|---|---|---|---|
| 6 | SF6_ENCODE_predictions_merge_metadata_2025-02_freeze1_98cols.csv.xz | Results section 3 | Fig. 3A<br>Supp. Fig. 6B-C<br>Supp. Fig. 8A-B,D | Fully trained classifier predictions generated using ENCODE data for the following classification tasks:<br><br>- Experiment assay , 11 classes: 7 ChIP + 2 RNA protocols + 2 WGBS protocols<br>- Experiment assay, 7 classes: IHEC core ChIP + input<br>- Cancer status, 2 classes<br>- Biomaterial type, 4 classes<br>- Donor sex, 3 classes<br>- Biospecimen source, 16 classes<br>- Donor life stage, 5 classes<br><br>Also includes metadata fields downloaded directly from the ENCODE Data Portal.<br>Capital-letter suffixes indicate the corresponding ENCODE metadata group: FILE, BIOSAMPLE, BIOSAMPLE_TYPE, and EXPERIMENT. |
| 7 | SF7_ChIP-Atlas_predictions_20240606_merge_metadata_freeze1_63cols.csv.xz | Results section 3 | Fig. 3B-F<br>Supp. Fig. 6D | Fully trained classifier predictions generated using ChIP-Atlas data for the following classification tasks:<br><br>- Experiment assay, 7 classes: IHEC core ChIP + input<br>- Cancer status, 2 classes<br>- Biomaterial type, 4 classes<br>- Donor sex, 3 classes<br>- Donor life stage, 5 classes<br><br>Also includes metadata fields downloaded directly from the ChIP-Atlas and GEO data portal.<br>The "DOWNLOADED" refers to ChIP-Atlas source information, per opposition to programmatically "extracted" fields from those. |

| | | | |
|---|---|---|---|
| 8 | SF8_imputation_effect_predictions.<br>csv.xz | Results section 3 Supp. Fig. 9D | Assay predictions of the two 6-class core-histone-mark assay classifiers (trained pval ChIP-seq tracks) used to assess the impact of training on imputed data. M-obs (model-observed) was trained on observed EpiATLAS tracks, and M-imp (model-imputed) on EpiATLAS-imputed tracks.<br><br>Comet training detail experiment keys:<br>Model-observed: dc1276654755410f89d87a804d8d68d9<br>Model-imputed: bce1ec47bdc0426ca4f38ad7b9eb77c5<br><br>One row per sample, from three inference sets:<br>- EpiATLAS imputed (n=9,570): predicted by M-obs only.<br>- EpiATLAS observed pval ChIP-seq (n=5,337): predicted by M-imp only.<br>- ChIP-Atlas (n=27,978): predicted by both models. This is a subset of the Fig. 3B file set.<br><br>By construction, the M-obs columns are empty for observed EpiATLAS files and the M-imp columns are empty for imputed EpiATLAS files, because these are the models' own training files.<br><br>Column details:<br>- ID: EpiATLAS md5sum (observed), full imputed track filename (imputed), or ChIP-Atlas experiment accession.<br>- EpiRR_no-v: EpiATLAS epigenome identifier.<br>- inference_data: EpiATLAS imputed / EpiATLAS observed / ChIP-Atlas.<br>- Expected assay: assay used as ground truth. It is the EpiATLAS assay metadata, or the ChIP-Atlas expected assay derived from the consensus of the database annotations.<br>- Predicted assay M-obs / predScore M-obs: predicted class and its probability (maximum softmax output) for M-obs.<br>- Predicted assay M-imp / predScore M-imp: same for M-imp. |

| | | | | |
|---|---|---|---|---|
| 9 | SF9_recount3_merged_preds_metadata_freeze1_43cols.csv.xz | Results section 3 | Fig. 3G-J<br>Supp. Fig. 6E | Fully trained classifier predictions generated using recount3 data for the following classification tasks:<br><br>- Experiment assay , 11 classes: 7 ChIP + 2 RNA protocols + 2 WGBS protocols<br>- Cancer status, 2 classes<br>- Biomaterial type, 4 classes<br>- Donor sex, 3 classes<br>- Donor life stage, 5 classes<br><br>Also includes metadata fields downloaded directly from the recount3 data portal. |
| 10 | SF10_rna_read_handling_predictions.csv.xz | Results section 3 | Supp. Fig. 5G | EpiATLAS 10-fold cross-validation predicted class labels and associated scores for RNA-Seq samples, comparing three read-handling variants of the same experiments:<br>- Unique: multi-mapped reads excluded (signal used for classifier training)<br>- UniqueMultiple: multi-mapped reads included<br>- Unstranded: the two Unique stranded tracks (plus and minus) summed into one signal per experiment ('signal_id' is an internal identifier and 'strand' is empty; use epirr to match with the other variants<br><br>Predictions are given for validation-fold samples only, generated by the fold classifiers for the following classification tasks:<br>- Experiment assay, 11 classes: 7 ChIP + 2 RNA protocols + 2 WGBS protocols<br>- Cancer status, 2 classes<br>- Biomaterial type, 4 classes<br>- Donor sex, 3 classes<br>- Biospecimen source, 16 classes<br>- Donor life stage, 5 classes<br><br>Each row includes the EpiRR identifier, strand, signal identifier, expected label, predicted label, and prediction score.<br>The classifier is identified by its experiment key, which can be cross-referenced with the model table. |

| | | | | |
|---|---|---|---|---|
| 11 | SF11_epirr_grouping_10fold_CV_preds.zip | Methods | NA | Two related zipped files.<br><br>EpiATLAS 10-fold cross-validation predicted class labels and associated scores of the MLP experiment assay (11 classes: 7 ChIP + 2 RNA protocols + 2 WGBS protocols) and biospecimen source (16 classes) classifiers, trained with two cross-validation grouping strategies (see Methods):<br><br>- UUID: all track types of a dataset are assigned to the same fold (grouping used throughout the paper).<br>- EpiRR: all datasets of an epigenome are assigned to the same fold. No epigenome has files in more than one validation split. One row per file and grouping strategy.<br><br>To reproduce the Supplementary Table 18 metrics for the Assay classifiers, merge the RNA (mrna_seq, rna_seq into rna_seq) and WGBS (wgbs-standard, wgbs-pbat into wgbs) protocols by summing their scores, take the class with the highest merged score as the predicted class, then compute the metrics per grouping strategy and split. |
| 12 | SF12_IHEC_ChromGene_assts--col1-5.tsv | Methods | Supp. Fig. 2 | Alternative sets of genomic coordinates used as input features for classifier training: protein-coding genes |
| 13 | SF13_StackedChromHMM_EnhancerMaxK27acCorrelations_n303114.tsv.xz | Methods | Supp. Fig. 2 | Alternative sets of genomic coordinates used as input features for classifier training: H3K27ac and gene correlation from ChromHMM |
| 14 | SF14_cpg_regions_topvariance_n303114.tsv.xz | Methods | Supp. Fig. 2 | Alternative sets of genomic coordinates used as input features for classifier training: 200 bp regions with most variable methylation |
